# The lipid-mediated mechanism of mechanosensitive channel MscS inactivation

**DOI:** 10.1101/2024.01.22.576751

**Authors:** Elissa Moller, Madolyn Britt, Fei Zhou, Hyojik Yang, Andriy Anishkin, Robert Ernst, Juan M. Vanegas, Sergei Sukharev, Doreen Matthies

## Abstract

Interpretations of experimental conformations of mechanosensitive channels gated by ‘force from lipids’ become more reliable when native lipids are preserved in the structures. *Escherichia coli* MscS is an adaptive osmolyte release valve that regulates turgor in osmotically stressed cells. MscS promptly opens under abrupt super-threshold tensions in the membrane, but at lower and more gradually applied tensions, it silently inactivates from the closed state. A central question has been whether to assign the commonly observed non-conductive conformation with splayed peripheral helices to a closed or inactivated state. We present a 3-Å MscS cryo-EM structure obtained in Glyco-DIBMA polymers, which avoid complete lipid removal. Within the complex, we observe densities for endogenous phospholipids intercalating between the peripheral and pore-lining helices. The lipidomic analysis shows a 2-3 fold enrichment of phosphatidylglycerol in Glyco-DIBMA-extracted MscS samples. The computed pressure of these lipids on the protein surface enforces the characteristic kinks in the pore-lining helices, sterically stabilizing the separation of the peripheral helices. Mutations of residues coordinating lipids in the crevices eliminate inactivation, allowing us to classify this group of structures as the inactivated state. Our study reveals a novel inactivation mechanism in which intercalated lipids physically decouple the tension-sensing helices from the gate.

## Introduction

MscS is a ubiquitous prokaryotic mechanosensitive channel of small conductance first identified in *Escherichia coli* ^1^ where it acts as an adaptive low-tension-threshold osmolyte release valve that regulates turgor pressure and ensures osmotic shock survival. It is the key cytoplasmic membrane component that provides bacterial adaptation to freshwater, which is the main transmission route for commensals and pathogens ^2^. The unique structural design of MscS was evolutionarily preserved and diversified in several paralogs present in the genomes of free-living bacteria ^1,3^. Multiple MscS homologs are found in all clades of walled organisms, including archaea, protists, fungi, and plants ^4–6^. In eukaryotes, MscS homologs fulfill sensory functions ^7,8^ and impart osmotic stability to internal organelles ^9,10^. The MscS structure’s manifestation in multimodal adaptive behavior warrants detailed analysis.

*E. coli* MscS is a homo-heptamer with each 287-residue protomer comprising three transmembrane helices and forming a large hollow cytoplasmic domain as revealed by the first crystal structure (PDB 2OAU ^11,12^). MscS is gated directly by tension transmitted through the surrounding lipid bilayer ^13,14^. Patch-clamp experiments in spheroplasts show that MscS readily opens in response to abruptly applied super-threshold tension but gradually inactivates under prolonged application of near-threshold tension. Its functional cycle comprises three functional states and two alternative transitions: closed-to-open and closed-to-inactivated, the latter state being non-conductive and tension-insensitive ^15–17^. The opening-closing transitions are sharply tension-dependent, essentially non-dissipative, and can be very fast ^17–20^. In contrast, the inactivation and recovery transitions are slow ^16,17^ but can be modulated by depolarizing voltages ^19^ and crowding pressure in the cytoplasm ^21^. Analysis of mutations shows that the propensity toward inactivation depends on the flexibility of pore-lining helices ^15^, salt bridges between the TM and cytoplasmic domains ^20^, hydrophobicity of TM2-TM3 crevices ^22^, and the residues that anchor the protein to the inner leaflet ^19^. While MscS opens robustly in most tested lipid environments ^13,14,23,24^, inactivation is not observed when MscS is reconstituted in soybean lipids ^13,14^, suggesting that lipids may be inseparable players in the mechanism of MscS inactivation and maintenance of the active state.

Although there are three functional states, only two classes of MscS structures exist. The first crystal structure of *E. coli* MscS, solved in Fos-choline-14 (PDB 2OAU ^11,12^), is to the first from the class characterized by splayed peripheral helices. The TM1-TM2 helical pairs in these conformations are disconnected from the sharply kinked TM3s that form the transmembrane pore and the gate. The hydrophobic gate of the 2OAU structure was found de-solvated in MD simulations ^25^, and thus the splayed structure was deemed non-conductive. More recent cryo-EM structures solved in mixed micelles or lipid nanodiscs (PDBs 6UZH, 7ONL, 7OO6, 7OO8, 6LRD, 6PWN, 6PWP, 6YVK ^26–30^) are more complete but generally retrace the original 2OAU ^11,12^ conformation. The second type of MscS conformation was observed in the A106V mutant (PDB 2VV5 ^31^), adopting an expanded barrel structure. This conformation was classified as open. However, estimations suggest that this conformation only achieves half of the experimental conductance of the fully open MscS (See Extended Data Fig. 1 and Text). Soon after, a similar expanded structure of WT MscS was solved in detergent DDM ^32^ and more recently in nanodiscs formed of short-tail lipids ^28,33^. These semi-open conformations have never been observed in the presence of lipids with regular full-length tails. The semi-open conformation of WT MscS, which is stable in a pure detergent environment (2VV5 class), can be converted into a splayed non-conductive state (2OAU class) simply by adding a low mole fraction of regular phospholipids ^26^. Notably, the two observed classes of conformations do not explain the entire three-state functional cycle of MscS.

Non-conductive splayed conformations of MscS have highly developed lipid-facing surfaces with characteristic crevices. Higher resolution structures detected densities of lipids stably associated with the splayed non-conductive structure of MscS, with one lipid consistently found closer to the periplasmic surface and more lipids on the cytoplasmic side ^26–29^. Simulations have shown that the splayed non-conductive structures specifically produce substantial distortion of the lipid bilayer on the cytoplasmic side, pulling the lipids into the crevices below the bilayer boundary and orienting them parallel to the bilayer plane ^33,34^. In addition, the fluorescence of tryptophan substitutions in the crevices detected an exchange of brominated lipids between these regions and the bulk bilayer ^34^. The splayed lipid-filled structure was interpreted as the closed state, and the hypothetical re-distribution of lipids between the crevices and the bilayer, driven by membrane tension, has been proposed as the opening mechanism ^35^. A subsequent finding of single-tail lipid-like densities inside the pore of MscS reconstituted into nanodiscs prompted an even more elaborate model of MscS gating by the ‘hook lipid’ and single-tail lipids traveling in and out of the pore ^27^.

Presenting the MscS gating cycle as two states (closed ↔ open) ^27,33,35^ is a simplification that neglects the second non-conductive functional state of MscS, the prominent inactivated state ^16,17^, which holds clear biological importance ^36,37^. To address this gap, we determined the structure of MscS extracted by the novel Glyco-DIBMA polymer ^38^, which preserves the positions of native lipids associated with the complex. During a typical membrane protein purification, lipids are largely removed by detergents prior to structural determination (Extended Data Fig. 2b). Detergents mimic the lipid bilayer environment to some extent but can cause structural changes ^39^. After solubilization, exogenous lipids are typically reintroduced in the form of mixed micelles, liposomes, or synthetic nanodiscs (Extended Data Fig. 2c-e) ^39^. However, extraction from the native membrane with detergent is still a prerequisite step. A more recent approach is to use amphipathic polymers to extract membrane proteins directly out of the lipid bilayer (Extended Data Fig. 2f), avoiding a complete de-lipidation step ^40^. Retaining a part of the local lipid environment provides the necessary detail on what lipids copurify and in what specific positions they associate with the protein ^41–46^.

In this work, we resolve the MscS conformation to 3 Å with intercalated lipids in specific conformations in the crevices and determine the key residues stabilizing their positions. We use Mass Spectrometry (MS) to determine the lipid species that preferentially associate with the complex. Molecular Dynamics (MD) simulations show the stable residence of these non-bilayer lipids that physically separate the lipid-facing helices from the pore-lining helices that form the gate. Computations of local 3D pressure exerted by the intercalating aliphatic tails reveal the pattern that stabilizes the characteristic kinked conformation of the pore-forming helices. We then mutate the residues that coordinate the lipid headgroups and analyze the mutant’s functional behavior by patch-clamp electrophysiology. Our data indicate that the splayed structure with intercalated lipids represents the inactivated state, which illustrates a novel lipid-mediated inactivation mechanism that uncouples the tension-receiving peripheral helices from the gate.

## Results

### Cryo-EM of MscS in native nanodiscs

Out of the three polymers we attempted (see Extended Data), only Glyco-DIBMA ^38^, a novel polymer that has not been used previously for cryo-EM, produced the whole *E. coli* MscS structure. Upon extraction and cobalt affinity column purification, the identity of polymer-extracted MscS was confirmed by SDS-PAGE and Western Blot (Extended Data Fig. 3d,g). Despite the high fraction of heavy particles inside large membrane patches, we proceeded to cryo-EM based on the 2D class averages from negative staining EM data collection (Extended Data Fig. 3m,p). The resulting map of MscS extracted with Glyco-DIBMA reached an average resolution of 2.99 Å, with the cytoplasmic domain estimated at 2.6 Å (Fig. 1). The structure is non-conductive, with the TM helices tilted inward at the top and splayed away from the pore-lining helix. The 2D class averages that contributed to this 3D reconstruction comprised the isolated particles associated with small lipid patches around the protein.

**Fig. 1:**
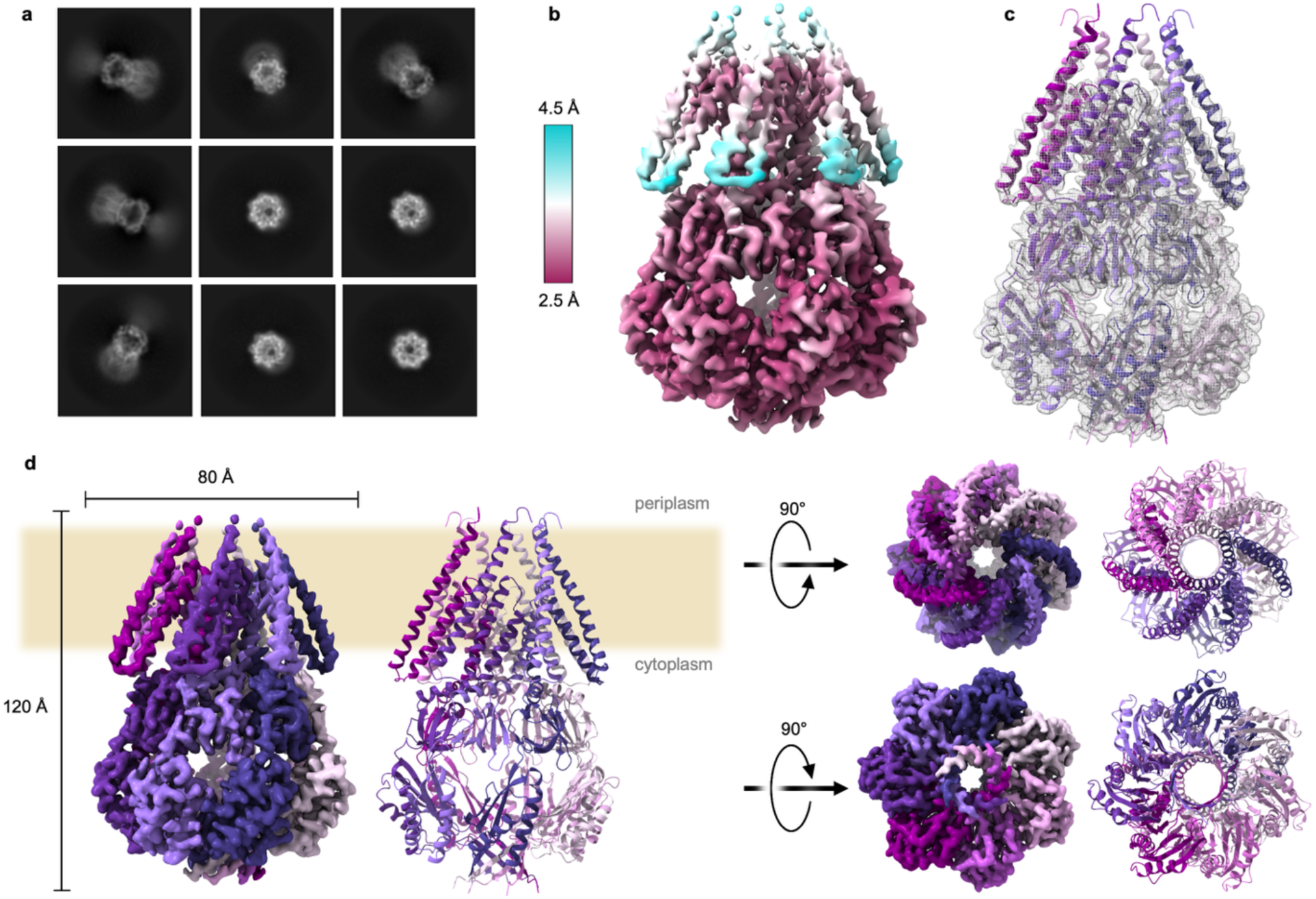
*E. coli* MscS cryo-EM structure in Glyco-DIBMA. (a) Selected 2D class averages of the particles used for the final reconstruction with a box size of 360 px (∼300 Å). (b) Cryo-EM map filtered and colored by local resolution. (c) Fitted model in ribbon representation inside map shown in gray mesh. (d) Map and model colored by chain shown as side view (left), top, and bottom view (right).

Densities unaccounted for by the protein model were examined for the presence of characteristic aliphatic chains and modeled as four individual phospholipids designated as Lipids 1 to 4. Because the resolution of dynamic lipids is insufficient to discern the headgroups, both POPE and POPG were possible candidates. Guided by lipidomic MS data (below), we were able to infer the lipid species and simulate their dynamic behavior. However, to avoid overinterpretation, in the final structure, the lipids are modeled with the headgroup truncated after the phosphate. The aliphatic tails of Lipid 1 (Fig. 2a) are oriented along TM1 and TM2 helices of neighboring chains, with the phosphate group latching to R88, forming a characteristic ‘hook’ ^27^. Lipids 2-4 (Fig. 2b-d) are oriented with the tails parallel to the inner surface of the membrane bilayer, wedged between the TM1-2 helices of neighboring chains. The protein-lipid contacts are stabilized by the electrostatic interactions between the lipid headgroups and charged residues in the gaps and crevices, most noticeably R88 for Lipid 1 and R59, K60, and D67 for Lipids 2-4. Multiple nonpolar contacts between hydrophobic residues and the aliphatic chains also occur. A full list of nearby residues stabilizing lipids in the static cryo-EM density-based structure is given in Extended Data Table 1. The extraction by the Glyco-DIBMA polymer ^38^ produced lipid patches of various sizes around the protein. The electron density surrounding the small native nanodisc patches from which we could resolve the structure is tighter than is typically seen in a pure detergent micelle. This polymer coat, along with the other remaining unmodeled densities that were not clearly defined lipids or proteins, is shown in Extended Data Fig. 4a,b. As was reported previously for detergent-extracted MscS reconstituted in synthetic lipid nanodiscs ^27^, we also observe several lipid-like densities inside the hydrophobic section of the channel pore. These particles do not have densities that could represent a phospholipid headgroup and appear to represent single aliphatic chains. As will be discussed below, these are possible *E. coli* fatty acid metabolites that are produced in the cytoplasm ^47^ and quickly exported to the periplasm ^48^.

**Fig. 2:**
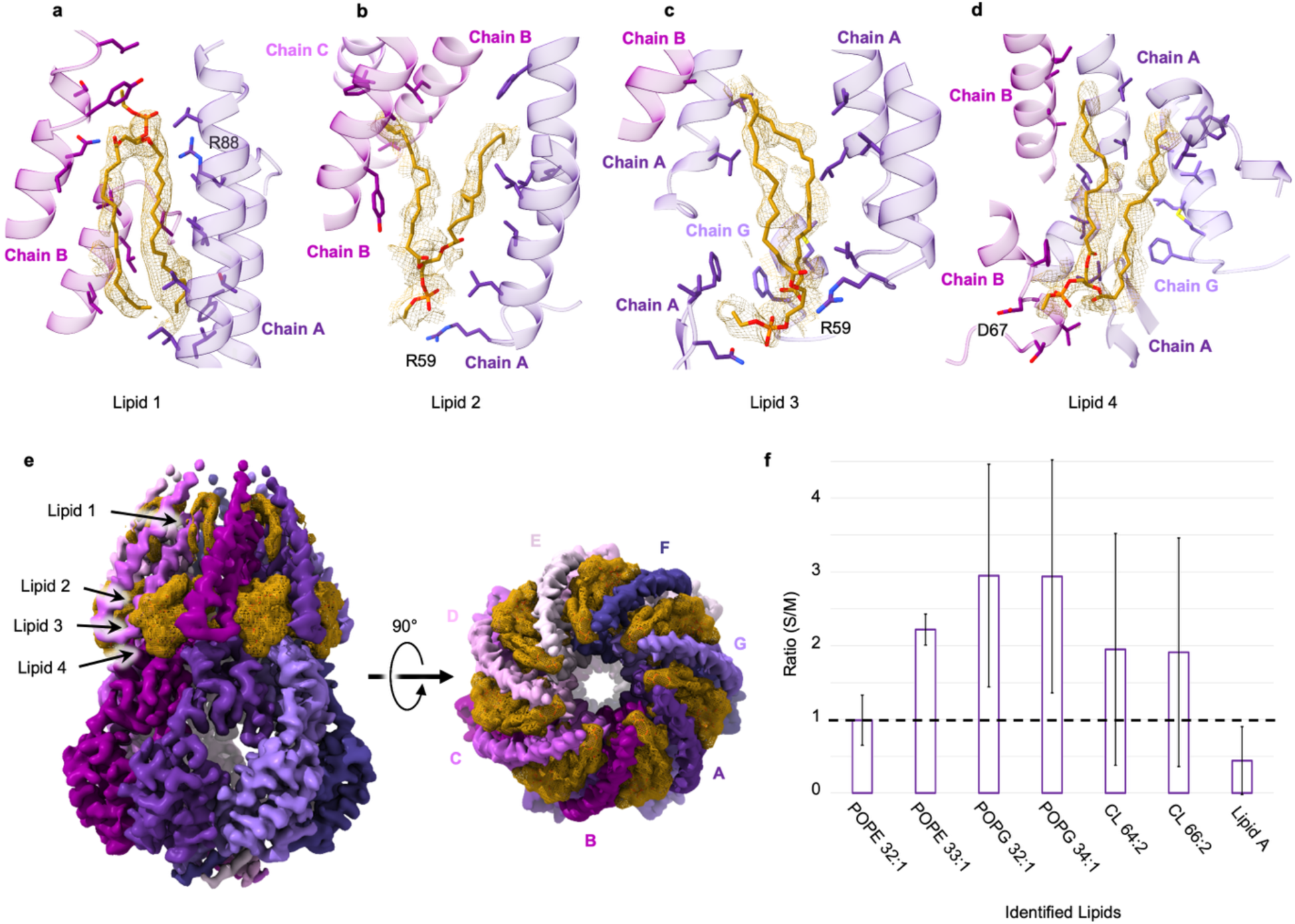
Resolved lipids in the TM region of *E. coli* MscS and lipid mass spectrometry analysis. Lipid 1 (a), Lipid 2 (b), Lipid 3 (c) and Lipid 4 (d) in yellow mesh density shown with all residues within 5 Å within the modeled protein. A full list of nearby residues is given in Extended Data Table 1. (e) Full map colored by chain with lipid densities shown in yellow mesh, side and top view. (f) Histogram comparing the relative amounts of lipids between the protein sample isolated in native nanodiscs (S) and the total inner membrane fraction (M). The ratio of mole fractions of specific lipids in these two samples (S/M) shows that POPE 33:1 and the two POPGs are enriched (1.5- to 2-fold), whereas other lipids, including POPE 32:1, are not. Cardiolipins are slightly enriched, whereas lipid A is depleted. Error shown is standard deviation. Exact values from this histogram are given in Extended Data Table 2.

### Mass Spectrometry of MscS-copurified lipids

MALDI-QTOF MS analysis was conducted on lipids copurified with MscS during extraction with Glyco-DIBMA and enrichment on a cobalt affinity column. The affinity-purified particles were subjected to an additional step of gel-filtration chromatography to isolate the smallest particles containing primarily annular lipids closely associated with the protein (Extended Data Fig. 4f). Fig. 2f and Extended Data Table 2 provide a comparison of the lipids co-purified with MscS with the composition of the inner membrane fraction isolated from the same expresser strain using a sucrose gradient. Using the Fast Lipid Analysis Technique (FLAT) -MS method for the characterization of lipid A and phospholipids ^49^, two highly abundant POPE species with one unsaturated bond, two prevalent POPG species, and two cardiolipin (CL) species were identified (Extended Data Fig. 4g). The *E. coli*-specific lipid A shows its typical m/z of 1796.21. A table comparing the relative amounts of lipids between the two samples, inner membrane (M) and protein (S), shows that odd chain POPE 33:1 and the two POPGs are enriched approximately 2-fold, other lipids POPE 32:1, Cardiolipins, are slightly enriched, whereas lipid A is largely excluded from the protein-associated pool (Fig. 2f).

### Lipid coordination and dynamics in the crevices

To better understand the dynamics of protein-lipid interactions, we embedded the solved structure with lipids in a POPE/POPG (3:1) lipid bilayer, equilibrated the system with soft restraints on the protein backbone for 20 ns, and ran for 100 ns in unrestrained simulations. Based on the lipid enrichment seen in MS, we embedded POPG as Lipids 1-3 and POPE as Lipid 4. The representative snapshots of lipid conformations in interhelical crevices are shown in Fig. 3, where Lipids 1-4 are color-coded individually. While the overall positions of lipids remained stable, they engaged with a broader set of surrounding polar sidechains compared to the static cryo-EM structure. In these simulations, Lipid 1 maintains a stable contact with R88 at the periplasmic side of MscS, and it is the only membrane lipid that contacts this residue. The average total probability of the headgroup contact is about 1.5 per subunit, half of which engage with the fatty acid oxygens of Lipid 1, and the other half with the phosphate group. Lipids 2, 3, and 4 interact primarily with the residues R59 and D67 located at the cytoplasmic side. Fig. 3b-d show that the aliphatic chains of the intercalated lipids frequently go around the TM2 helix and thus physically separate it from the TM3s forming the hydrophobic gate (L105-L109). Besides that, these lipids form electrostatic contacts with R74. The contact probabilities were scored for each individual lipid, emphasizing electrostatic interactions of phosphate and fatty acid oxygens or amino groups with neighboring sidechains. The statistics of polar contacts within 5 Å of each specific group averaged over all subunits during the last 80 ns period of the simulation are shown in Extended Data Table 3. Fig. 3c-d show the multiplicity of H-bonding patterns between the headgroups within the three-lipid cluster, illustrating the low entropic restraint allowing for a stable residence of three lipids in the crevice in multiple configurations.

**Fig. 3:**
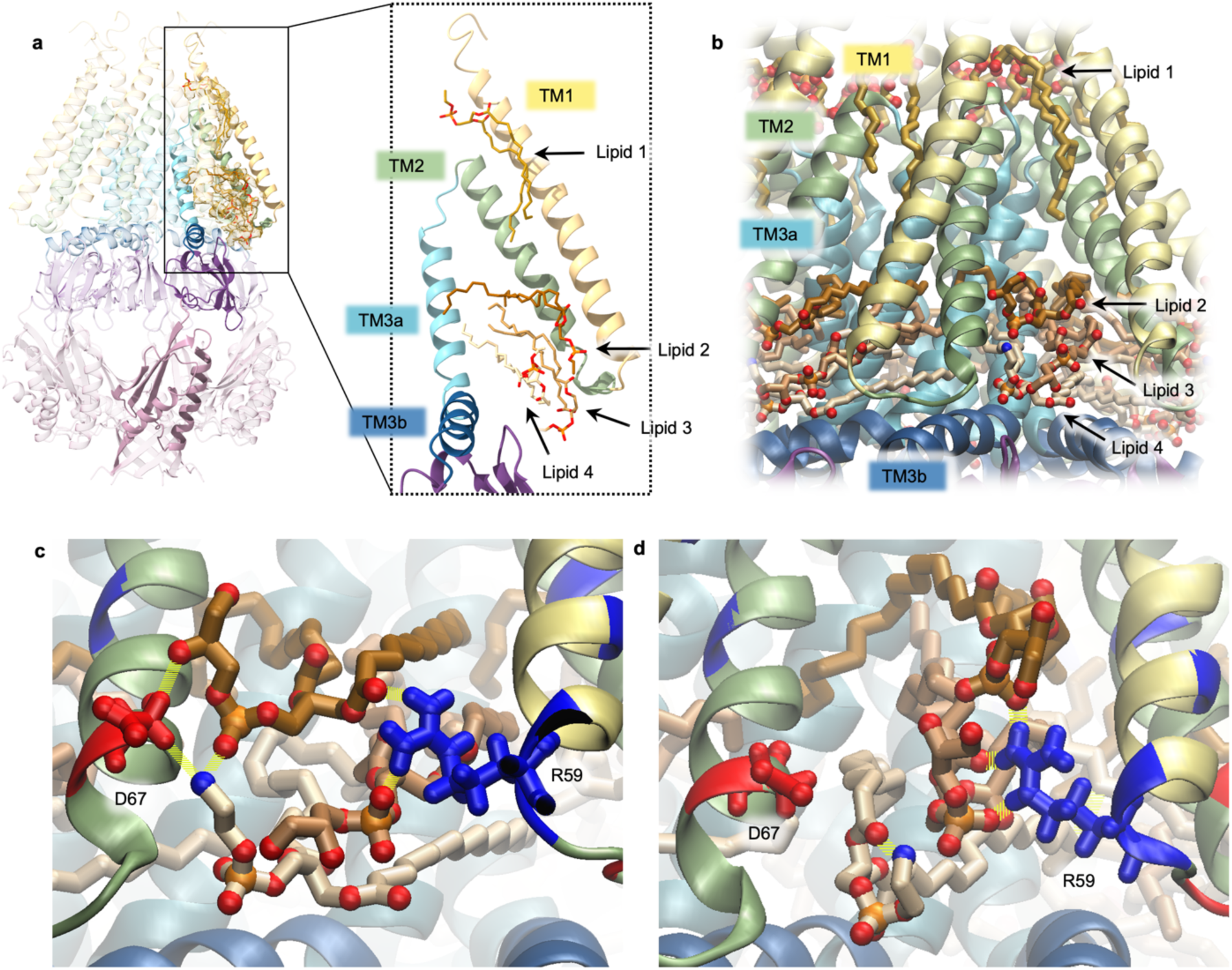
Molecular dynamics simulations of lipids in MscS’s interhelical crevices. (a) The model of MscS built into the cryo-EM density with the resolved lipids, colored by domain with TM1 in yellow, TM2 in green, TM3a in cyan, and TM3b in blue. (b) A snapshot from MD simulations of the modeled lipids showing lipid tail penetration between TM1-TM2 pairs (yellow and green) and pore-forming TM3a (cyan). (c-d) Zoomed-in snapshots illustrating two different lipid clusters stabilized by inter-lipid H-bonds and coordinated between D67 and R59. The clusters appear to be both flexible and stable due to their multitude of configurations. Lipids are shaded from dark to light: Lipid 2 (dark), Lipid 3 (medium), and Lipid 4 (light). The statistics of intercalated lipid contacts in interhelical crevices with neighboring residues are listed in Extended Data Table 3.

Fig. 4 shows the local stress distribution on the surface of individual MscS chains through the traction vector. The 3D local stress was computed from a parallel simulation of the solved MscS structure in GROMACS and analyzed with the GROMACS-LS package (see Methods). The normal component of the traction shows various ‘hot spots’ of pushing (blue) and pulling (red) forces on the surface of the channel subunits (Fig. 4b), which incorporates interactions with the surrounding lipids and solvent molecules. The enlarged region near the partially de-solvated portion of the pore (Fig. 4c) shows pulling forces, while prominent pushing forces exerted by lipid chains are concentrated precisely in the kinked region of TM3 near L111 and G113. The pressure of intercalated lipids, therefore, stabilizes the G113 kink previously associated with the inactivated state ^15^.

**Fig. 4:**
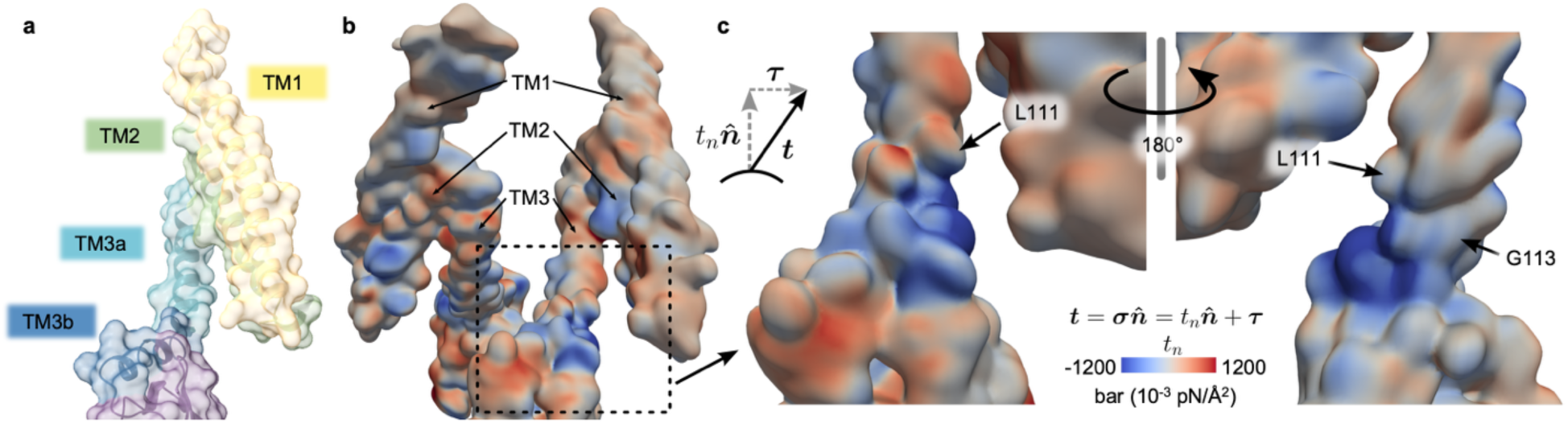
Computation of local 3D stress acting on the protein surface of the lipid-embedded MscS. Stress on the surface of individual MscS chains is visualized through the traction vector, 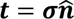, which is obtained from the product of the local stress tensor (***σ***) and the normal unit vector (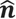) defined by the protein surface. (a) MscS surface colored by domain. (b) Normal component of the traction 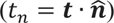 on chains A (left) and E (right) of the simulation showing the distribution of forces on TM1, TM2, and TM3. (c) Close-up of chain E in the vicinity of the kink in TM3 near residues L111 and G113. Blue colors indicate forces pushing into the surface, while red colors indicate pulling forces. The surface of the MscS chains is computed as isosurfaces from the average mass density of the protein.

### Functional effects of crevice mutations

To investigate how protein interactions with the intercalating lipids affect the function of MscS, we conducted functional tests of several mutants using patch-clamp (Fig. 5). The ‘hook’ Lipid 1 is observed in many structures ^26–29^. It was predicted to play a role in the gating transitions by detaching and moving to the bilayer during the opening transition ^27^. Fig. 5a-b shows patch-clamp traces comparing WT MscS and R88A, illustrating the depth of inactivation and the kinetics of recovery. The small hydrophobic substitution R88A had a mild phenotype, with no significant changes to activating tension (Extended Data Table 4) or percent inactivation but did show a subtle slowing down of the recovery from inactivation (Extended Data Table 5). In our cryo-EM samples, the densities of aliphatic chains of Lipid 1 are clear, but the nature of the headgroup cannot be easily identified. The interactions of the guanidine group of R88 with the phosphate would stabilize almost any type of lipid in this place.

**Fig. 5:**
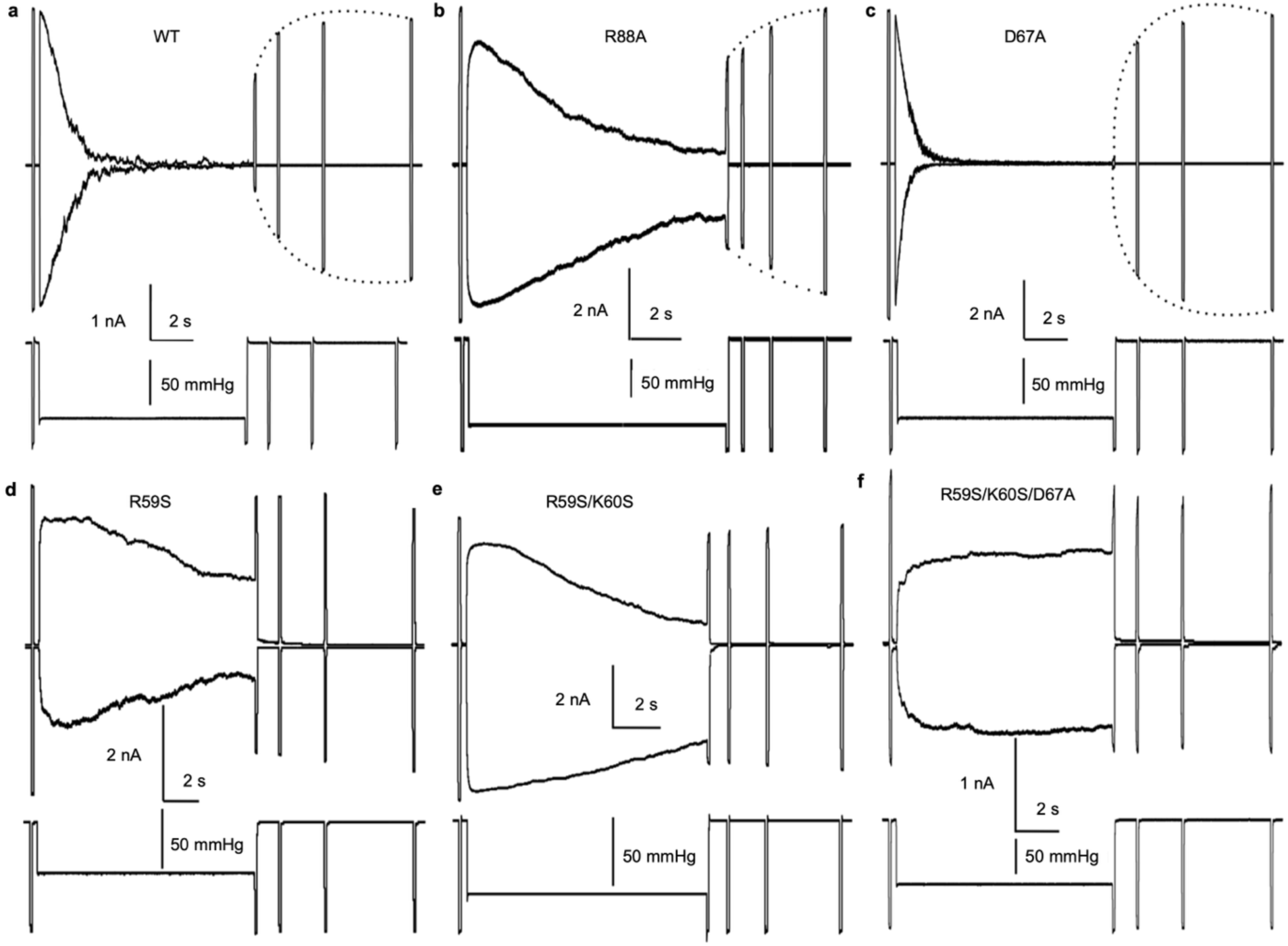
Patch-clamp traces examining inactivation and recovery. The first short test pulse of saturating amplitude opens the entire population of channels, then the extended step of intermediate tension drives them toward inactivation. The following train of saturating test pulses monitors the fraction of remaining active channels, which is used to determine the kinetics of recovery. R88A (b) behaves similarly to WT (a). The D67A mutant (c) inactivates completely and recovers very quickly, whereas R59S (d) is mainly resistant to inactivation. The double mutant (e) and triple mutant (f) follow the R59S Inactivation/Recovery phenotype. The triple mutant is also resistant to adaptive closing at the midpoint tension.

The polar headgroups of Lipid 2 and Lipid 3 predominantly interact with R59’s sidechain, perhaps with additional contribution from K60. Lipid 4 intermittently interacts with the nearby D67. We conducted recordings using the inactivation/recovery protocol after introducing point mutations at R59, D67, and K60 (Fig. 5). All single substitutions for R59 resulted in a non-inactivating phenotype (Extended Data Table 5). R59S has the least amount of inactivation at +30 mV pipette potential (Fig. 5d), the voltage at which WT is more resistant to inactivation. The traces show no sign of recovery, suggesting that this mutant greatly resists inactivation under these conditions. Conversely, D67A inactivates completely, recovers very quickly when the pipette pressure drops to zero (Fig. 5c), and causes a slight increase in activation midpoint (Extended Data Table 4). Substitutions at K60 have an intermediate phenotype, with slightly more inactivation and a slower rate of recovery (Extended Data Table 5). Double R59S/K60S and triple R59S/K60S/D67A are both strongly non-inactivating (Fig. 5e,f). Based on these results, we conclude that the lipid intercalated state of the TM2-TM3 crevice, stabilized primarily by R59 interacting with the lipid phosphates and D67 interacting with the amino groups of POPE, corresponds to the inactivated state of the channel. The lipid penetration is ‘invited’ by R59, and when the lipids get into the crevice, they are additionally stabilized by D67. It is important to note that in the D67A mutant, R59 is in place and electrostatically unopposed due to the absence of the nearby D67 chain. Therefore, D67A is more willing to inactivate, but the lack of a counterpart for interaction with the amino group of POPE (Lipid 4) may make POPE leave the crevice more quickly, and so the recovery is quick. The lipid expulsion from the crevices might be cooperative, which is why the departure of Lipid 4 may promote the exit of other lipids, resulting in the compaction of the transmembrane domain and the re-establishment of the functionally predicted TM2-TM3 contact ^22^.

### Comparison of structures and lipid positions

The backbone of the Glyco-DIBMA polymer-extracted MscS is most similar to PDB 6PWN ^27^ solved in synthetic nanodiscs, and shares the same architecture as the 2OAU-class with the TM1-2 helices splayed away from the pore-lining helix (Extended Data Fig. 5a). The lipid-coordinating side chains are clearly resolved, and the modeled lipid positions are the most similar to PDB 6LRD ^29^ (Extended Data Fig. 5h-j), but unlike those exogenously added synthetic lipids, endogenous *E. coli* inner membrane lipids are retained here. In semi-open DDM and LMNG solubilized structures, detergent molecules replace these lipids on the periphery of the protein (PDBs 7OO0, 7ONJ ^26^). Without this de-lipidation caused by detergent, a new lipid (Lipid 2) was also resolved.

PDB 6PWN ^27^ has additionally modeled 7 single lipid tails without density for lipid headgroups in the pore of the protein above the gate oriented along TM3a. We also see disordered density in these regions but did not model them as lipids as they are along the symmetry axis of the protein and lack clear densities for headgroups (Extended Data Fig. 4a,b). Detergents have been modeled in these locations previously ^26^, but since we never used detergent in our preparation, our densities likely correspond to free fatty acids which are necessary intermediates in phospholipid metabolism ^47,50^. The Glyco-DIBMA polymer could have ‘mobilized’ fatty acids during the extraction and let them penetrate the pore. Moreover, these free fatty acids are actively extruded to the extracellular space ^48^. MS shows a number of molecular species smaller than phospholipids co-purified with the protein sample (See Extended Data Fig. 4g), but at this point, we cannot identify these species unequivocally. Based on the essentially frictionless opening process ^51^ and the microsecond kinetics of MscS closing ^19^, we have no reason to connect the presence of these particles in the pore with the gating process. More likely, the gating transitions reflect tension-dependent wetting and de-wetting of the hydrophobic gate ^25^.

## Discussion

The results presented above answer the central question: Does the splayed non-conductive conformation represent the closed or inactivated state? We provide evidence that MscS harboring lipids between the splayed peripheral helical pairs is in the inactivated state. The solved structure had sufficient resolution to identify residues coordinating lipids between the helices (R59 and D67), and mutations of these residues destabilizing lipids in the crevices directly affected inactivation. The analysis of lipids tightly associated with Glyco-DIBMA extracted complexes showed some enrichment of anionic lipids, presumably residing in the crevices. The MD simulations of the solved structure illustrate how the aliphatic tails of lipids in the crevices penetrate between the TM2 and TM3 helices, separate the membrane-facing (TM1-TM2) helical pairs from the gate, and exert pressure that stabilizes the characteristic G113 kink in TM3, previously associated with the inactivated state ^15^. The conclusion that the splayed state represents the inactivated state implies the existence of the alternative non-conductive conformation with TM1-TM2 pairs reconnected with the TM3 barrel (‘compact’ closed state) and is highly consistent with the three-state ‘dashpot’ gating mechanism of MscS ^16^. This conclusion fully corroborates the dramatic functional effects of hydrophilic substitutions in the crevices, which lead to immediate tension-driven channel inactivation without opening ^22^. It also explains the nature of the two alternative conformational pathways, fast opening and slow inactivation, both originating from the closed state ^17–19^.

The choice of the novel Glyco-DIBMA polymer ^38^ advanced our characterization of MscS, revealing the positions of native lipids. Instead of complete de-lipidation, the polymer ‘cuts’ the protein out of its native membrane together with associated lipids, which could be resolved in their native positions. In the course of extraction and purification, no exogenous lipids were added, and detergents were never used. The overall resolution of the protein was near 3 Å (Fig. 1), and the densities of four protein-intercalating lipids (per subunit) were resolved (Fig. 2). The parallel MS analysis indicated POPG enrichment. The positions and dynamics of POPG/POPE lipids in the crevices were tested in MD simulations of the solved structure, which confirmed stable coordination of Lipid 1 by R88 and preferential coordination of the cytoplasmic three-lipid clusters (Lipids 2-4) by R59 and D67. The lipid motions inside the cytoplasmic side clusters were substantial as H-bonds interlinked the lipid headgroups in many different configurations (Fig. 3), indicating minimal entropic penalty and favorably low energy.

Mutations of lipid-coordinating residues had different functional manifestations. R88 coordinating Lipid 1 was previously proposed to play a role in gating by liberating this hook lipid during the opening transition 1. ^27^. Our patch-clamp results, in contrast, show that R88A substitution does not affect the opening, inactivation, or recovery processes substantially (Fig. 5). However, the effects of uncharged substitutions for R59 and D67 were more dramatic. The R59S mutant showed a vivid non-inactivating phenotype, indicating that destabilization/depopulation of the cytoplasmic lipid cluster in the crevice prevents inactivation. The single D67A substitution exhibited fast inactivation and fast recovery, suggesting that R59 alone is able to stabilize lipids in the crevice and lead to inactivation, but in the absence of the second ‘attachment point’ for the lipid cluster, the lipid exchange between the bilayer and the crevice becomes faster. The triple R59S/K60S/D67A mutation that removes most of the polar attachment point for lipids prevents inactivation. This result indicates that the lipid-intercalated structure represents not the closed but the inactivated state. The functional behavior of these mutants alongside the lipid-filled structure allows us to make this assignment for the entire class of ‘splayed’ structures.

Previous studies have already attempted to rationalize and interpret the initial crystal structure of MscS solved in a splayed conformation with sharply kinked TM3 helices (PDB 2OAU ^11,12^). The hydrophobic gate in this structure is de-solvated, and the conformation must be non-conductive ^25^. Soon after, our functional study of MscS revealed the presence of the prominent tension-insensitive inactivated state ^16^. In patch-clamp, the channel population opened fully in response to an abrupt tension application, but only a fraction of channels responded to a slow tension ramp of the same amplitude, while the majority silently transitioned to the inactivated state. This behavior, which allows MscS to discriminate the stimuli according to their dynamics, was coined as the “dashpot” tension coupling mechanism. It was proposed that the splayed crystal structure must represent the inactivated state ^16^. Indeed, the PDB 2OAU ^11,12^ conformation shows the peripheral lipid-facing helices (TM1-TM2) disconnected from TM3 that form the gate. The sharp kink at G113 orients the TM3b helix almost parallel to the membrane interface and prevents the splayed TM1-TM2 pairs from closing the gap and straightening along TM3s. The G113A mutation imposing higher helical propensity to the kink region was shown to suppress inactivation ^15^. This result suggested that the G113 kink is not present in all conformations, and there must be a compact state with the TM3 gate region reconnected with peripheral helices through a predicted hydrophobic ‘clutch’ formed by the rings of conserved hydrophobic residues, F68 and L72 on TM2 and L111 and L115 on TM3 ^22^. The “dashpot” was predicted to transmit the abruptly applied tension from TM2 to TM3 and open the gate. Under slowly applied tension, the “dashpot” was predicted to ‘give up’ and mechanically uncouple the peripheral helices from the gate. We now believe that the intercalating lipids resolved in the crevices act precisely as a viscous dashpot filler mediating a slow/delayed uncoupling. Moreover, our MD results show that the tails of crevice-intercalated lipids exert pressure on the G113 kink region (Fig. 4), thus stabilizing this kink that keeps the TM1-TM2 pairs separated from the gate.

What conformations may explain the three-state kinetic scheme with two independent opening and inactivation transitions? Fig. 6 shows the place of the solved structure in the three-state functional cycle of MscS. The structure represents a stable inactivated state. The expulsion of lipids from the crevices is predicted to compact the structure into a narrowly packed closed state. In this state, the peripheral (TM1-TM2) helical pairs form a hydrophobic contact with TM3s. At the same time, the characteristic kink at G113 relocates to conserved G121 to allow for compaction ^15^. The reconnected TM3 and TM2 helices provide the transmission route for the tension received from the lipids by the peripheral TM1-TM2 pairs, producing the conformation that is now capable of opening by tension. From that state, the channel may undergo two alternative tension-driven transitions, opening or inactivation. The opening of MscS, a steeply tension-dependent process, is predicted to occur through tilting and complete straightening of pore-lining TM3 helices, which is accompanied by a 16-18 nm^2^ in-plane protein expansion. Opening is extremely fast and was shown to be essentially non-dissipative ^51^. Inactivation is associated with a smaller effective expansion, and it is shown to be a much slower process dependent on the nature of lipid-anchoring residues ^19^. This is highly consistent with the slow penetration of lipids into the crevice, thus literally implementing a ‘dashpot’ mechanism.

**Fig. 6:**
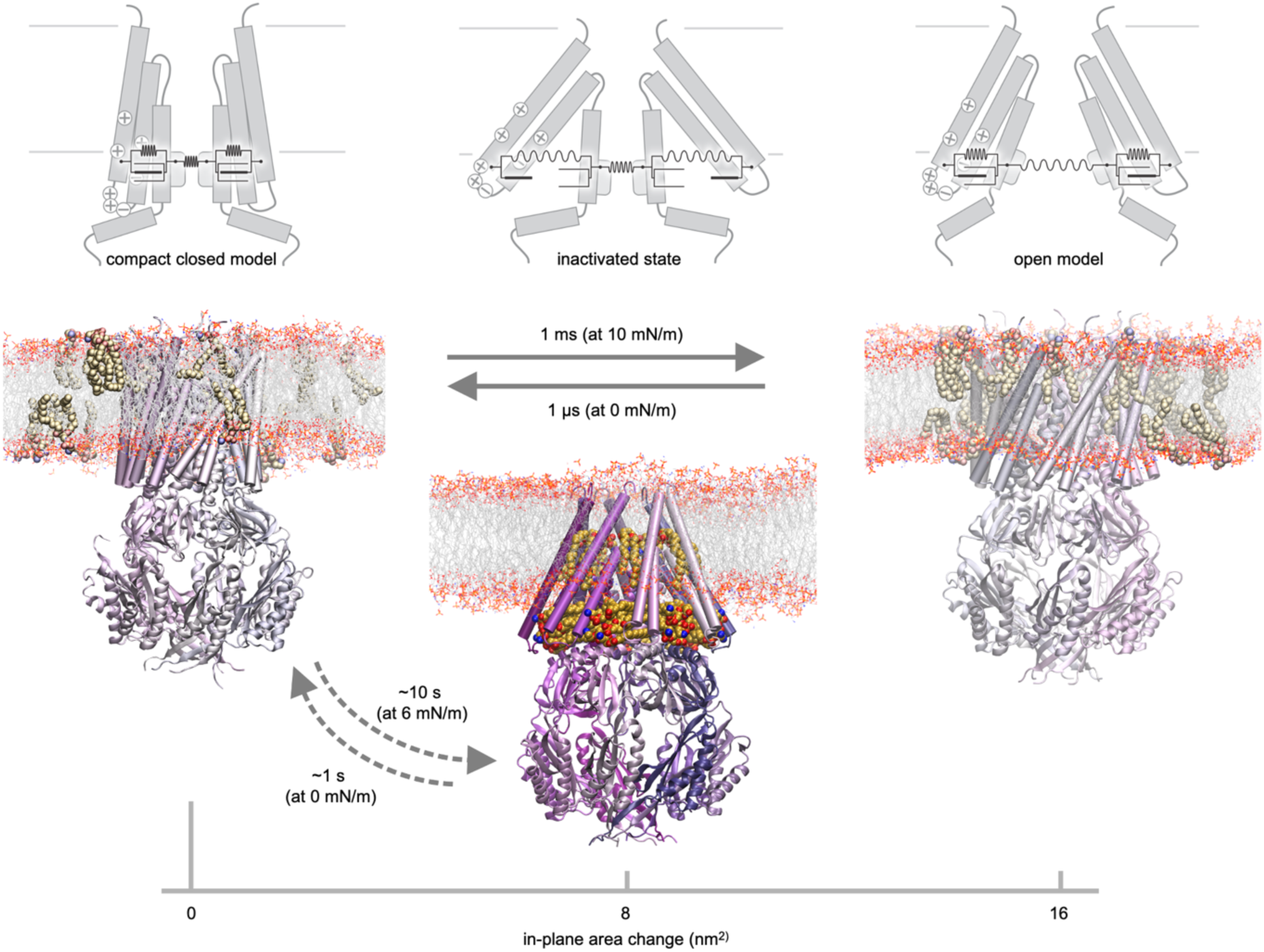
The experimentally determined kinetic scheme of MscS and the assignment of the newly solved structure. The opening-closing and inactivation-recovery pathways are characterized by different spatial scales and drastically different characteristic times of transitions. The solved cryo-EM structure, filled with native lipids, fits all the criteria of the inactivated state. The transition to the inactivated state from the closed state is accompanied by moderate in-plane protein expansion and occurs at near-threshold tensions. The opening from the hypothetical compact closed state is associated with a larger lateral expansion at higher tensions. The gating scheme implies a sharp kinetic separation between the opening and inactivation pathways.

We could not find any evidence in the literature that the splayed conformation is definitively the closed state. Repeated attempts to open the splayed conformation in simulations were generally unsuccessful ^33,34,52^. The presentation of MscS as a two-state (closed-open) system without inactivation found in these papers is an oversimplification that overlooks the third functional state. Not only is the inactivated state seen in patch-clamp traces, but its functional importance has also been demonstrated in osmotic viability experiments ^36,37^. What is the functional role of the inactivated state? The integrity of the cytoplasmic membrane is critical for bacterial energetics ^53,54^. Inactivation is a safeguard against spurious channel opening and leaky state of the membrane in actively metabolizing, hyperpolarized, and turgid cells. Transition into the inactivated state occurs when there is a need (1) to reduce the active population and limit the permeability response to low osmotic shocks to minimize metabolic losses, (2) to properly terminate the full permeability response to a strong down-shock involving both MscS and MscL ^36,37^ and (3) to eliminate metabolite leakage from a swollen cell when the peptidoglycan is compromised by either β-lactam antibiotics or lysozyme. The splayed lipid-stabilized inactivated state appears to be the most stable (default) state, preventing leakage. When the channel is isolated in the presence of regular lipids, this state is the only conformation that has been experimentally observed. The assignment of the splayed lipid-filled conformation to the inactivated state now puts all of the previous functional results and modeling predictions ^15^ into place: the G113 kink previously associated with the inactivated state to be stabilized by intercalating lipids; straightening this kink would allow the TM2 to reconnect with TM3 to form a hydrophobic ‘clutch’ for force transmission from the periphery to the gate ^22^.

In conclusion, this work shows that applied to the MscS channel, the ‘force-from-lipids’ gating mechanism is twofold: lipids at the protein boundary transmit external forces to the gating apparatus to open the channels, but at the same time, protein-penetrating lipids can interrupt the force transmission route and make the channel tension-insensitive. The presented findings give the first explanation of the extremely broad range of MscS’s kinetic responses, from millisecond opening rates to tens of seconds for inactivation. The opening transition can be very fast and essentially frictionless, as it does not seem to involve deep penetration of lipids. The slow dynamics of lipid penetration into secluded locations in the protein, leading to inactivation, kinetically separates the two independent expansion pathways of opening and inactivation, which imitates a dashpot. The inactivated state is most stable, whereas the compact closed conformation appears unstable in isolation, and the fully open state is transient and requires tension. For these reasons, the closed and fully open states have not yet been visualized by structural methods. Special means will be required to stabilize these conformations and solve them experimentally.

## Methods

### MscS Expression and Inner Membrane Isolation

*E. coli* MscS wild-type (WT) was expressed in MJF465 cells (Frag1 ΔmscL::Cm, ΔyggB, ΔmscK::kan) ^1^ from the pB10d vector, a modified pB10b plasmid ^55^. The plasmid is Isopropyl β-D-thiogalactopyranoside (IPTG) inducible with the lac UV5 and lacIq promoters, confers ampicillin resistance, and includes a C-terminal His6-tag on the MscS gene. It uses the high-copy-number ColE1/pMB1/pBR322/pUC origin of replication and the T7 bacteriophage ribosome binding site. The MJF465 strain originated from the Ian Booth laboratory (University of Aberdeen, Aberdeen, UK) was provided by Dr. Samantha Miller (University of Aberdeen). A single colony of the plasmid-containing strain was inoculated into 35 mL Lysogeny Broth (LB) media supplemented with 100 µg/mL Ampicillin (Amp) and grown overnight. 5 mL of the overnight culture was sub-cultured into six 2 L flasks containing 1 L LB supplemented with Amp and grown for 2 hours shaking at 225 rpm at 30°C to an OD600 of approximately 0.3 before induction with 1 mM IPTG for an additional 3 hours to a final OD600 of 2.0. When the target OD was achieved, cells were placed on ice to prevent further growth, pelleted at 7,000 rpm using Sorvall GSA rotor for 15 min at 4°C, and resuspended in French Press Buffer (20 mM Tris/HCl pH 7.6, 150 mM NaCl) supplemented with 200 µL saturated PMSF in EtOH to 30 mL total volume before rupturing with a French press. Unbroken cells were pelleted by centrifugation at 10,000 rpm using Eppendorf F-34-6-38 rotor for 15 min at 4°C, and the supernatant was then ultracentrifuged at 25,000 rpm using Beckman SW28 rotor (112,400 x g) for 1 hour at 4°C to pellet membranes. Supernatant was discarded, and membranes were resuspended in 20% [w:v] sucrose prepared in Gradient Buffer (20 mM HEPES pH 7.2, 5 mM EDTA) and homogenized before being placed on top of a 3-step sucrose gradient of 0.77 M, 1.44 M, and 2.02 M prepared in Gradient Buffer, and finally ultracentrifuged at 25,000 rpm using Beckman SW28 rotor (112,400 x g) for 16 hours at 8°C. The inner membranes were harvested from the upper band of the gradient by puncturing the tube wall with a syringe needle, diluted in Gradient Buffer with no sucrose to 70 mL total volume, and ultracentrifuged at 25,000 rpm using Beckman SW28 rotor (112,400 x g) for 1 hour at 4°C to re-pellet membranes. The membrane pellets were flash-frozen in liquid nitrogen and stored at −80°C until use.

### Polymer Extraction and Affinity Purification

*E. coli* inner membranes were resuspended to 10 mg/ml in Membrane Buffer (20 mM HEPES pH 7.4, 100 mM NaCl) and homogenized before the addition of 0.5% [w:v] of solid Glyco-DIBMA ^38^ (Cube Biotech). The first condition (Glyco-DIBMA #1) was allowed to extract overnight at 4°C on the tube revolver, while the second condition (Glyco-DIBMA #2) was allowed to extract for 2 hrs at room temperature on the tube revolver. The insoluble fraction was then pelleted by ultracentrifugation at 25,000 rpm using Beckman SW28 rotor (112,400 x g) for 35 min at 4°C. The supernatant was then diluted 1:5 with Membrane Buffer to improve binding efficiency and incubated overnight on the tube revolver at 4°C with TALON Cobalt Resin (Takara Bio). The next day the resin was washed with 10 column volumes of Membrane Buffer, 10 column volumes of Wash Buffer (20 mM HEPES pH 7.4, 100 mM NaCl, 10% glycerol [v:v]) with 10 mM imidazole, 5 column volumes Wash Buffer with 20 mM imidazole, and 5 column volumes of Wash Buffer with 40 mM imidazole before elution with Elution Buffer (20 mM HEPES pH 7.4, 100 mM NaCl, 10% glycerol [v:v], 350 mM imidazole). Elution fractions were then dialyzed with Membrane Buffer overnight to remove imidazole and glycerol and concentrated the following day using a 100 kDa ultrafiltration membrane (Amicon) to the desired concentration for cryo-EM grid preparation. Protein fractions were resolved and analyzed using SDS-PAGE, Western Blot, and BN-PAGE (see Extended Data Methods); gel images and Western blots can be found in Extended Data Fig. 3. Many different polymers are now commercially available; we have also attempted MscS extraction using SMALP 200 ^43^ and CyclAPol C_8_-C_0_-50 ^44^, but these preparations did not allow us to resolve the transmembrane domain of MscS (see Extended Data Text).

### Negative Staining Electron Microscopy

3 µL of MscS sample at 0.3 mg/ml (30 times higher than MscS in detergents or mixed micelles of typically ∼ 0.01 mg/mL) was applied to a glow-discharged carbon coated 400 square mesh copper grid (CF400-CU, EMS) and incubated for 1 min at room temperature. The grid was then blotted by filter paper (Whatman Grade 2) and washed once with 3 µL of Nano-W negative staining solution (Nanoprobes) followed by incubation with 3 µL of Nano-W for 1 min. The grid was blotted to remove excess staining solution and air dried by waving for an additional minute. Images were recorded using an FEI Tecnai T20 TEM operated at 200 kV with a direct electron detector K2 Summit (Gatan Inc). Data was collected using SerialEM ^56^ at a nominal magnification of 25,000x with a pixel size of 1.479 Å/px, a dose rate of approximately 7 e^−^/px/s with 0.2 s frames over 10 s, and a defocus range between −1.3 and −2 µm. A total of 651 images were collected, and Bsoft 2.2.0 ^57^ was used to unpack the image stacks. Image processing was performed using cisTEM v1.0.0. ^58^. 37,902 particles were picked from the micrographs after CTF estimation ^59^ using a maximum radius of 60 Å and a characteristic radius of 40 Å with a threshold of 5. Particles were extracted with a box size of 168 px (∼250 Å) and analyzed using 2D classification (Extended Data Fig. 3p-r). 2D class averages and data collection parameters for MscS extracted with SMALP 200 ^43^ and CyclAPol C_8_-C_0_-50 ^44^ can be found in Extended Data. Negative staining micrographs for the polymers alone in buffer (see Extended Data Methods) are illustrated in Extended Data Fig. 6.

### Cryo-EM Grid Preparation and Data Collection

3 μL of purified MscS between 0.1 and 0.7 mg/mL was applied to a glow-discharged (15 s at 15 mA using EasyGlow) Quantifoil R 1.2/1.3 + 2 nm C 400 Cu mesh grid (EMS), a holey grid with a thin 2 nm continuous carbon film. The grids were blotted for 6 s at 4°C and 95% humidity and plunged frozen into liquid ethane using a Leica EM GP2 (Leica) and stored in liquid nitrogen. For the Glyco-DIBMA extracted MscS, a double sample application was performed where 3 µL of the sample was applied, blotted by hand using Whatman Grade 2 filter paper, then 3 µL was re-applied and blotted using a Leica EM GP2. The grids were screened on an FEI Tecnai T20 TEM before data collection. Cryo-EM datasets were acquired with SerialEM ^56^ using a G1 or G4 Titan Krios (FEI, now ThermoFisher Scientific) operated at 300 kV and equipped with an energy filter and K3 camera (Gatan Inc.). Movies of 50-60 frames were collected with a total dose of 50 e^−^/Å^2^ at a nominal magnification of 105,000x corresponding to a physical pixel size of 0.83 or 0.8469 Å/px at a dose rate of 9-15 e^−^/px/s and a defocus range of −0.8 to −1.8 µm. Specific collection parameters for each dataset can be found in Extended Data Table 6.

### Cryo-EM Data Processing

The specific workflows for image processing are illustrated in Extended Data Fig. 7-11. All processing was performed with cryoSPARC v3.3.250 ^60^. Movies were processed with patch motion correction ^61^ and patch CTF estimation ^59^. Blob picker was used for the majority of particle picking, and particles were extracted with a box size of 320-360 px (∼300 Å) and subjected to 2D classification to remove junk particles. Selected particles were used for ab initio reconstruction, heterogeneous refinement, and non-uniform refinement using no symmetry (C1) as well as 7-fold symmetry (C7) ^62^. The local resolution was estimated, and the maps were filtered accordingly. Additional CTF refinements and local motion correction did not improve resolution, most likely because the 2 nm carbon layer support already reduces beam-induced motion ^63^.

### Model Building and Refinement

For the MscS atomic model, PDB 7OO6 ^26^ was rigid body-fitted into the sharpened local resolution filtered map using UCSF ChimeraX v.1.8 ^64^. The model was then manually rebuilt in COOT v.0.9.8.92 ^65^ using the same map, which was generated from non-uniform refinement with C7 applied. The truncated lipid was generated using the SMILES string CCCCCCCCCCCCCCCC(=O)OCC(COP([O-])(=O)OCC)OC(=O)CCCCCCC\C=C\CCCCCCCC in eLBOW ^66^. Iterative rounds of manual refinement in COOT and real space refinement in Phenix v.1.21.2-5419 ^67^ were performed. The quality of the model and fit to the density was assessed using MolProbity ^68^ and Phenix ^67^. Side chain densities for a helix and beta sheet are shown in Extended Data Fig. 12. In contrast to the structured helix reported in 6PWN ^27^ and 6YVK ^28^, the N-terminal residues are poorly resolved until residue 17, indicating that they are flexible and disordered. The protein was modeled from residue 17 to 281 out of the 287 total residues (Fig. 1c), with the side chains removed from residues 17, 18, 20, 21, 280, and 281 due to low density. All structural figures were prepared using UCSF ChimeraX v.1.8 ^64^. Cryo-EM map and model analysis values are listed in Extended Data Table 7. PDBs have been made available together with the cryo-EM maps with the following EMD/PDB IDs: EMD-71088/7P0N for the MscS/Glyco-DIBMA (C7) and EMD-71089/7P0O for the MscS/Glyco-DIBMA (C1). Movies as well as particle stacks have been uploaded to EMPIAR ^69^.

### FLAT-MS for lipid analysis

The MscS sample for Mass Spectrometry (MS) lipid analysis to identify co-purified lipids was prepared as described for Glyco-DIBMA #2, with an additional gel filtration step using a hand-packed column containing Sephacryl S-200 Superfine (Pharmacia Fine Chemicals). No dialysis was performed, the sample was exchanged into Membrane Buffer (20 mM HEPES pH 7.4, 100 mM NaCl) on the column. The SDS-PAGE, Western Blot, and Negative Staining micrographs from this purification showing the removal of large lipid patches with size exclusion can be found in Extended Data Fig. 4c-f. The control sample is purified *E. coli* inner membranes (see *MscS Expression and Inner Membrane Isolation)* resuspended in Membrane Buffer. An initial round of MS without this additional separation step was done on the remaining protein sample after cryo-EM grid preparation, but it showed no dramatic lipid enrichment since the ratios were skewed by the large membrane sections rather than examining only the lipids immediately surrounding MscS (See Extended Data Fig. 4). 1 µL of sample pellet was scraped and plated directly onto a steel reusable MALDI plate. 1 µL of citric acid extraction buffer (0.2 M citric acid, 0.1 M trisodium citrate, pH 3.5) was spotted on top of the plated sample. The steel MALDI plate was placed into the chamber with water on the bottom and heated to 110°C in an oven for 30 minutes. After incubation, the plate was rinsed with water and air-dried, and 1 µL of norharmane matrix was spotted on each sample. A Bruker Matrix-Assisted Laser Desorption/Ionization trapped ion mobility spectrometry Time-of-Flight Mass Spectrometry MALDI (tims TOF) MS was used for the Fast Lipid Analysis Technique (FLAT), a protocol for direct MS biomass visualization of lipid A and phospholipids ^70^ as described below.

Our Bruker MALDI (tims TOF) spectrometer is equipped with a dual ESI/MALDI source with a SmartBeam3D 10 kHz frequency tripled Nd:YAG laser (355 nm); the system was operated in “qTOF” mode (TIMS deactivated) for these experiments. Ion transfer tuning was used with the following parameters: funnel1 RF, 440.0 Vpp; funnel2 RF,490.0 Vpp; multipole RF, 490.0 Vpp; CID energy, 0.0 eV; and deflection delta, −60.0 V. Quadrupole was used with the following values for MS mode: ion energy, 4.0 eV. The m/z scan range for MS is set to 1000 to 3000. Collision cell activation of ions used the following values for MS mode: collision energy, 9.0 eV; collision RF, 3900.0 Vpp. Agilent ESI tune mix was used to perform calibration of the m/z scale. MALDI parameters in qTOF were optimized to maximize intensity by tuning ion optics, laser intensity, and laser focus. All spectra were collected at a 104 μm laser diameter with a beam scan using 800 laser shots per spot and 70 and 80% laser power, respectively.

MS data were collected in negative ion mode. In all cases, 10 mg/mL of norharmane (NRM) in 1:2 MeOH/CHCl3 [v:v] was used for lipid detection. All MALDI (tims TOF) data were visualized using mMass (Ver 5.5.0). Peak picking was conducted in mMass using the following parameters: S/N threshold: 3.0; relative intensity threshold: 5.0%; picking height: 50; apply baseline and smoothing. For the triplicate experiment, we analyzed the samples on three different days, obtaining a total of nine mass spectra for both purified inner membrane and MscS protein samples. These spectra were analyzed using mMass to determine the average relative intensity values for each of the seven assigned lipids. Standard deviations for each lipid intensity were calculated using Microsoft Excel.

### Molecular Dynamics Simulations

The simulation system was assembled around our cryo-EM structure with three POPG and one POPE per subunit based on the enrichment observed in mass spectrometry. The lipid bilayer built with 498 POPE and 166 POPG molecules (ratio 3/1) was assembled around the channel to bring the lateral size of the orthogonal simulation cell to 150 x 150 Å, allowing for at least 9-10 lipid layers between the closest mirror images of the protein. The membrane protein system was hydrated, and K^+^ and Cl^−^ ions were added to make 200 mM KCl salt solution, bringing the total number of atoms in the system to ∼395,000. The simulation cell was assembled using CHARMM GUI ^71^. The entire system was energy minimized with a restrained protein backbone for 10,000 steps using the conjugate energy gradient algorithm and simulated for 20 ns with the protein backbone harmonically restrained near the modeled positions with a spring constant of 1 kcal/mol/Å. A flexible orthogonal cell was simulated with periodic boundary conditions under 1 bar pressure and a lateral tension of 20 dyne/cm. All simulations were performed using NAMD2 ^72^ with CHARMM27 force field ^73^, TIP3P water model, particle mesh Ewald method for long-range electrostatics estimation ^74^, a 10 Å cutoff for short range electrostatic and Van der Waals forces, and a Langevin thermostat set at 310 K. After the restrained backbone stage completed, the system was simulated unrestrained for 100 ns, with the last 80 ns used to quantify the contact statistics. Visualization of the channel structures was performed using Visual Molecular Dynamics (VMD) ^75^. All the structural and statistical analyses for MD simulations were performed using custom-written Tcl scripts for VMD. To compute the local stress/pressure in 3D, the equilibrated systems were converted to GROMACS ^76^ input files and simulated with version 2023 of the program for 50 ns, saving the positions and velocities to the trajectory every 5 ps. The saved trajectory was first re-centered around the center of mass of the protein to account for spontaneous lateral diffusion of the protein and then post-processed using the GROMACS-LS and MDStress library ^77,78^. This post-processing step produced a time-averaged 3D stress field for each system using a grid spacing of 0.2 nm and an electrostatics cutoff of 2.2 nm. The 3D local stress was then mapped onto the surface of the MscS individual chains by computing the traction vector (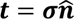) as the product of the stress tensor (***σ***) with the surface normal unit vector 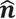 as previously reported ^79^. The protein surface and normal vectors were obtained by computing the isosurface of the 3D protein mass density. For ease of visualization, we only show the normal component of the traction vector 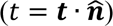. The traction data was visualized using the ParaView software ^80^.

### Mutagenesis and Patch-clamp Electrophysiology

Point mutations were introduced to the MscS plasmid using the Q5 site-directed mutagenesis kit (New England Biolabs). Giant spheroplasts were generated as described previously ^81^. In short, cells were grown in LB to turbidity and then sub-cultured 1:9 into a defined minimal LB at 250 mOsm and grown for 1.5 h in the presence of 0.06 mg/ml cephalexin, which prevents cell septation, before inducing with 1 mM IPTG for the final 30 min. The resulting filaments were transferred into 1 M sucrose with 1.8 mg/ml BSA buffer and treated with 0.17 mg/ml lysozyme in the presence of 4 mM EDTA, which degrades the peptidoglycan layer and causes the filaments to collapse into giant spheres. The reaction was stopped with 5 mM MgCl_2_. The giant spheroplasts were isolated via sedimentation through a one-step 1.2 M sucrose gradient at 2,000 rpm using Eppendorf Rotor A-4-81 for 5 min at 4°C.

Patch-clamp traces were recorded from giant bacterial cell spheroplasts. Population channel recordings were conducted at ±30 mV (pipette) on excised inside-out patches in symmetric buffer containing 400 mM sucrose, 200 mM KCl, 50 mM MgCl_2_, 5 mM CaCl_2_, and 5 mM HEPES at pH 7.4. Membrane patches were obtained using borosilicate glass pipettes (Drummond) and recorded on an Axopatch 200B amplifier (Molecular Devices). Negative pressure stimuli were applied using a high-speed pressure clamp apparatus (HSPC-1; ALA Scientific) programmed using the Clampex software (Molecular Devices). Pressure and voltage protocol programming and data acquisition were performed with the PClamp 10 suite (Axon Instruments). The patch responses were first recorded at +30 mV (pipette) with symmetric 1-second pressure ramps to determine the activation thresholds, midpoints, and saturation level for the plasmid expressed MscS population in both MJF465 (ΔMscL ΔMscS ΔMscK) and PB113 (ΔMscS ΔMscK) where MscL is also present for comparison. PB113 is derived from the MJF429 strain ^1^ with the additional ΔrecA ^82^ and was provided by Paul Blount (University of Texas at Dallas, Richardson, TX, USA). MscL can be used as an internal standard with a known midpoint tension γ_0.5_ = 14 mN/m and midpoint ratio of 0.6, meaning that the ratio of the tension and the pressure midpoints between expressed MscS and native MscL can be calculated ^83^. To observe the inactivation of MscS, an Inactivation/Recovery protocol was used ^16,17,19^: the full population of channels was opened with a saturating test pulse and then held at the midpoint tension for MscS for 10 s followed by three test pulses with zero pressure steps in between. The rate of recovery (τ) was determined by fitting the recovery pulses to a monoexponential function.

## Acknowledgements

We thank Patrick O’Reilly for composing and illustrating a detailed analysis of membrane protein extraction for Extended Data Fig. 2. We are grateful to Joshua Zimmerberg and Jennifer Peterson for access to the Tecnai T20 electron microscope, AJ Morton, Zabrina Lang, and Rick K. Huang for the support on the G1 Krios, and Wyatt Peele, Kedar Sharma, and Mario Borgnia for the support on the G4 Krios. We thank Samantha Miller for providing the MJF465 strain, Paul Blount for providing the PB113 strain, Manuela Zoonens for providing the CyclAPol C_8_-C_0_-50 polymer, Jake Rosetto for assistance with spheroplasting and patch-clamp, Gerald Kidd for assistance in membrane isolation, and Alex Sodt and Louis Tung Faat Lai for valuable discussions. The research was supported by the Division of Intramural Research of the *Eunice Kennedy Shriver* National Institute of Child Health and Human Development, NIH (grant NICHD intramural projects Z1A HD008998 to DM). This work was supported by NIH R01-AI135015 to SS. JMV acknowledges the support of the National Science Foundation through Grant No. CHE-1944892/2326678. This material is based upon work supported by the National Science Foundation Graduate Research Fellowship Program to EM under Grant No. DGE 1840340. Any opinions, findings, and conclusions or recommendations expressed in this material are those of the author(s) and do not necessarily reflect the views of the National Science Foundation. This work utilized the computational resources of the NIH HPC Biowulf cluster (http://hpc.nih.gov). Molecular graphics and analyses performed with UCSF ChimeraX, developed by the Resource for Biocomputing, Visualization, and Informatics at UCSF, with support from NIH R01-GM129325 and the Office of Cyber Infrastructure and Computational Biology, NIAID.

## Author Contributions

EM: membrane isolation, protein purification, biochemistry (SDS, Western, BN), negative stain microscopy and data collection, cryo-EM grid preparation, cryo-EM data processing, model building and refinement, mutagenesis, patch-clamp, data interpretation, figure making, writing, funding

MB: membrane isolation, patch-clamp, figure making, writing

FZ: biochemistry advice, cryo-EM grid screening, Krios data collection

HY: lipid mass spec, data interpretation, writing

AA: molecular dynamics simulations, data interpretation, writing

RE: lipid mass spec, data interpretation, writing

JMV: molecular dynamics simulations, stress and traction force computation, data interpretation, figure making, writing

SS: membrane isolation, sample preparation for mass spec, data interpretation, writing, supervision and training, funding

DM: model building, data interpretation, writing, supervision and training, funding

## Competing Interest Declaration

The authors declare no competing interests.

## Additional Information

Supplementary Information is available for this paper.

## Data Availability

The data that support this study are available from the corresponding authors upon request. Cryo-EM maps have been deposited in the Electron Microscopy Data Bank (EMDB) under accession codes EMD-71088 (C7 MscS in Glyco-DIBMA) and EMD-71089 (C1 MscS in Glyco-DIBMA). Raw movies will be uploaded to the Electron Microscopy Public Image Archive (EMPIAR). The atomic coordinates have been deposited in the Protein Data Bank (PDB) under accession codes 9P0N (C7 MscS in Glyco-DIBMA) and 9P0O (C1 MscS in Glyco-DIBMA). The source data for Extended Data Fig. 3 and 4 are provided in the Source Data file with this paper.

## Code Availability

The scripts used in this study are available from the corresponding authors upon request.

## Abbreviations

MscS: mechanosensitive channel of small conductance
TM: transmembrane
EM: Electron Microscopy
BN: Blue Native
MS: Mass Spectrometry
MD: Molecular Dynamics
POPE: phosphatidylethanolamine
POPG: phosphatidylglycerol
CL: cardiolipin

## Notes

### Competing Interest Statement

The authors have declared no competing interest.

### Summary of Updates

This is a complete revision of the previous version

