## Supplemental Data for "The lipid-mediated mechanism of mechanosensitive channel MscS inactivation"

**Extended Data**

Supplementary Text: Estimations of the conductance of the 2VV5 class of structures, *E. coli* MscS polymer extraction and comparison

Extended Data Fig. 1: Estimations of conductance.
Extended Data Fig. 2: Membrane protein extraction methods for structural studies.
Extended Data Fig. 3: Comparison of electron microscopy and biochemistry for the different polymers.
Extended Data Fig. 4: Modeled lipids, extra densities, and lipid mass spectrometry.
Extended Data Fig. 5: Structure comparison.
Extended Data Fig. 6: Negative staining EM of polymers.
Extended Data Fig. 7: Glyco-DIBMA #1 cryo-EM workflow.
Extended Data Fig. 8: Glyco-DIBMA #2 cryo-EM workflow.
Extended Data Fig. 9: SMALP 200 #1 cryo-EM workflow.
Extended Data Fig. 10: SMALP 200 #2 cryo-EM workflow.
Extended Data Fig. 11: CyclAPol $C_8$-$C_0$-50 cryo-EM workflow.
Extended Data Fig. 12: Quality of the cryo-EM map and fitted model of MscS in Glyco-DIBMA.

Extended Data Table 1: Residues within 5 Å of modeled lipids.
Extended Data Table 2: Values for lipid mass spectrometry analysis.
Extended Data Table 3: Statistics from MD simulations of lipid contacts.
Extended Data Table 4: Activation tension for mutants patched in pB113.
Extended Data Table 5: Summary of patch clamp data for inactivation and recovery.
Extended Data Table 6: Cryo-EM data collection parameters and analysis.
Extended Data Table 7: Cryo-EM map and model analysis.

Primer List

Supplementary Methods: Protein Validation using SDS-PAGE, Western Blot, and Blue Native-PAGE; SMALP 200 and CyclAPol $C_8$-$C_0$-50 Extraction and Affinity Purification; Negative Staining Electron Microscopy of Polymers; and Negative Staining Data Collection for SMALP 200 and CyclAPol $C_8$-$C_0$-50

Supplementary References

**Supplementary Text**

*Estimation of the pore conductance of the MscS crystal structure* (PDB 2VV5)

The 2VV5 structure [1] represents an expanded conformation of MscS stabilized by the A106V mutation. The A106 residue is located in the TM3a pore-lining helix inside the gate region formed by L105 and L109, which create a hydrophobic constriction. The A106's methyl sidechain faces the neighboring helix's G104 in a knob-in-a-hall manner, preserving tight helical packing. The bulkier V106 sidechains (in place of A106) push against the neighboring TM3, resulting in an expanded, presumably conductive state. A similar state of WT MscS was obtained in pure DDM (PDB 4HWA [2]) with no mutations. Because these two crystal structures were solved in a delipidated state without applied membrane tension, we believe they might be not expanded to their limit and therefore represent a sub-conductive (semi-open) state. Below, we present a 'macroscopic' conductance estimation of the 2VV5 conformation.

Extended Data Fig. 1a shows the top view of the 2VV5 pore, with the gate-keeping sidechains defining the minimal pore diameter. Next to it is the model of the fully open pore (opf), featuring completely straightened TM3 helices. This kink-free state of TM3 represents the structurally reasonable limit for pore expansion under tension, avoiding the disjoining of subunits. Below, side views of the two conformations are presented along with plots of average pore radii for each structure and variations in pore resistance along the z coordinate (pore axis). Pore radius was estimated as the surface-to-surface distance with the CHARMM36 version of VDW radii. Resistances were calculated in 5 Å slabs of aqueous solution inside the pore. All calculations were performed using custom-written Tcl scripts for VMD. The specific conductivity of the internal buffer is taken as 33 mS/cm, corresponding to the experimentally measured value for the standard patch-clamp buffer. Summing the resistances along the pore produces 0.49 nS single-channel conductance for 2VV5, which is half of the 1.1 nS for the opf model, which is the open-state model with completely straight TM3 helices. The 1.1 nS conductance corresponds to the experimental fully open MscS conductance in standard buffer. This relatively simple macroscopic estimation supports the conclusion that 2VV5 [1] and similar structures represent the semi-open conformation and should not be interpreted as the fully open state.

*E. coli MscS polymer extraction and comparison*

Three polymers were attempted in parallel under similar extraction conditions: SMALP 200 [3], CyclAPol $C_8$-$C_0$-50 [4] (commercially available as Ultrasolute Amphipol 18 from Cube), and Glyco-DIBMA [5]. SMALP was the original polymer that was shown to work for direct extraction and structural characterization, CyclAPols overcame many of the limitations associated with the SMALP chemistry including restrictions on pH and divalent cations, and Glyco-DIBMA is a novel polymer that has not been used previously for cryo-EM. For all three polymers the extraction was less efficient than detergent solubilization and proved to be most effective at room temperature rather than the typical 4°C. The concentrations of each polymer were chosen to be at the lowest end of the range given by the manufacturer. A larger range of concentrations from 0.1% to 2% were initially tested for SMALP 200, but decreasing the polymer concentration was detrimental to extraction efficiency and increasing it did not improve extraction in a linear fashion based on SDS-PAGE and Western Blots comparing the soluble and insoluble fractions after ultracentrifugation. As polymers have been previously shown to interact with membrane proteins the lowest effective range was chosen. Membrane concentrations from 10 mg/mL to 30 mg/mL were tested, and we found that increasing the dilution only marginally improved the extraction efficiency. Room temperature solubilization rather than at 4°C greatly improved extraction, likely due to the phase of the lipids at this temperature [6]. Higher temperatures have been used previously and proved effective [7] but were not attempted in this comparison. Even when allowed to extract overnight at 4°C the extraction was lower than at room temperature for 2 hours. Room temperature extraction for longer than 2 hours was not tested but may prove effective provided that the protein is stable. The buffers for SMALP 200 and CyclAPol $C_8$-$C_0$-50 were based on a previous publication [4]. HEPES was chosen for Glyco-DIBMA since the polymer is commercially available in that buffer. Negative Staining EM is shown for each of these polymers alone in the appropriate buffers at the concentrations used for the extractions in Extended Data Fig. 6. At medium magnification there is already a macroscopic pattern visible for each polymer, and at high magnification clusters of varying sizes are observed. Polymer extracted MscS was purified by a cobalt affinity column and analyzed by SDS-PAGE, Western Blot, and BN-PAGE (Extended Data Fig. 3d-l). SDS-PAGE and Western Blot confirmed the identity and purity of the sample. The BN-PAGE was inconclusive with the majority of the protein remaining in the well rather than running according to the size of the heptamer. BN-PAGE requires a higher protein concentration than SDS-PAGE, which may contribute to the lack of signal for the low yield polymer extracted sample. Despite this, the sample looked promising albeit heterogenous using Negative Staining EM with some isolated particles and clear MscS top views resembling rings even though an approximately 30 times higher protein concentration was necessary compared to detergent solubilized MscS or MscS in mixed micelles (Extended Data Fig. 3m-o). The SMALP 200 Negative Staining EM micrographs were the most similar in appearance to detergent with distinct particles and only some clusters/aggregation (Extended Data Fig. 3o). CyclAPol $C_8$-$C_0$-50 was noticeably more clustered/aggregated but still had distinct particles identifiable as MscS (Extended Data Fig. 3n). Glyco-DIBMA was extremely heterogeneous with entire membrane patches interspersed with the single particles (Extended Data Fig. 3m). For all three polymers the Negative Staining EM data collection and processing resulted in MscS-like 2D class averages (Extended Data Fig. 3p-r), and we proceeded to cryo-EM. The initial structural attempts to solve MscS in a native nanodisc with both SMALP 200 and CyclAPol $C_8$-$C_0$-50 were only partially successful under the purification conditions that were tested, resulting in side view 2D class averages with a blurry TM region and 3D maps with unresolved TMs. However, using the Glyco-DIBMA polymer to form native nanodiscs, yielded a complete 3D reconstruction. A selection of the 2D class averages used for the 3D reconstruction show clear features in the TM region in typical side view 2D class averages. These features are lacking in the blurry TM region from the SMALP 200 and CyclAPol $C_8$-$C_0$-50 extracted MscS averages (Extended Data Fig. 9-11). Another explanation could be that the binding of the other polymers to the lipid-protein complex is not stable enough and, as the buffers are lacking additional lipids and polymers to exchange with, they dissociate from the protein. If the polymers and lipids dissociate from the TM region those domains will become unstable and may start to disintegrate. It is important to consider however that these polymers were also tested in the presence of glycerol at a slightly different pH using Tris/HCl buffer instead of HEPES and may work well for structural studies under different conditions.

### Extended Data Figures

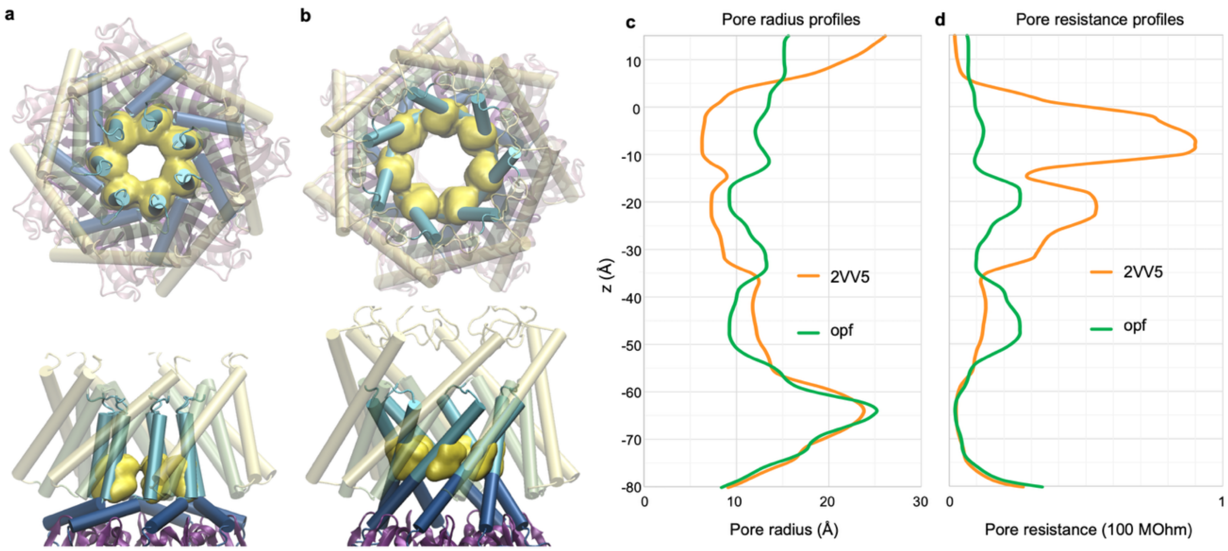


**Extended Data Fig. 1: Estimations of conductance for 2VV5 structure.** The two conductive conformations, 2VV5 [1] expanded state (a) and the modeled open state (opf) with kink-free pore-lining TM3 helices (b). The gate constriction is shown in yellow. The profiles of the pore radii (c) and resistance (d) for the two structures are plotted along the pore axis (z). The axis has its zero positioned at the midplane of the membrane. Integration of the resistance along the z-axis shows that 2VV5 has half the conductance of the fully open model (opf).

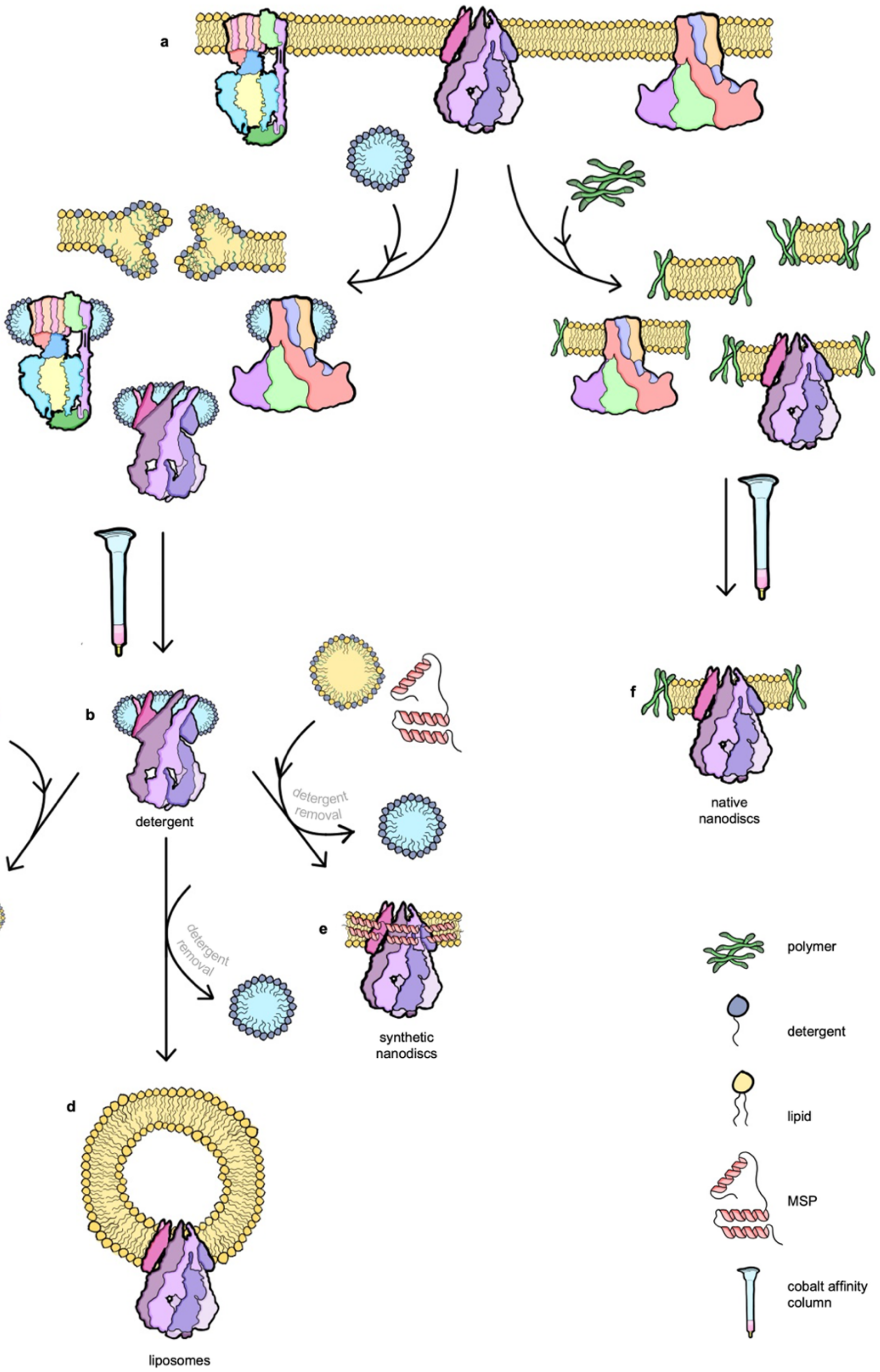


**Extended Data Fig. 2: Membrane protein extraction methods for structural studies.** (a) Membrane proteins in the bilayer [PDBs left to right 5ARE [8] (ATP synthase), 2OAU [9,10] (MscS), and 3JCF [11] (CorA)]. (b) Detergent solubilization leading to a conductive conformation of MscS (PDB 7OO0 [12]), followed by affinity chromatography. (c) Addition of detergent solubilized long chain lipids to form mixed micelles leading to a non-conductive conformation of MscS (PDB 7OO6 [12]). (d) Addition of detergent solubilized lipids followed by detergent removal for liposome reconstitution (e) Addition of detergent solubilized lipids and membrane scaffold protein (MSP) followed by detergent removal for synthetic nanodisc reconstitution leading to a non-conductive conformation of MscS (PDB 6LRD [13]). (f) Direct membrane protein extraction using polymers, followed by affinity chromatography. Illustrated by Patrick O'Reilly.

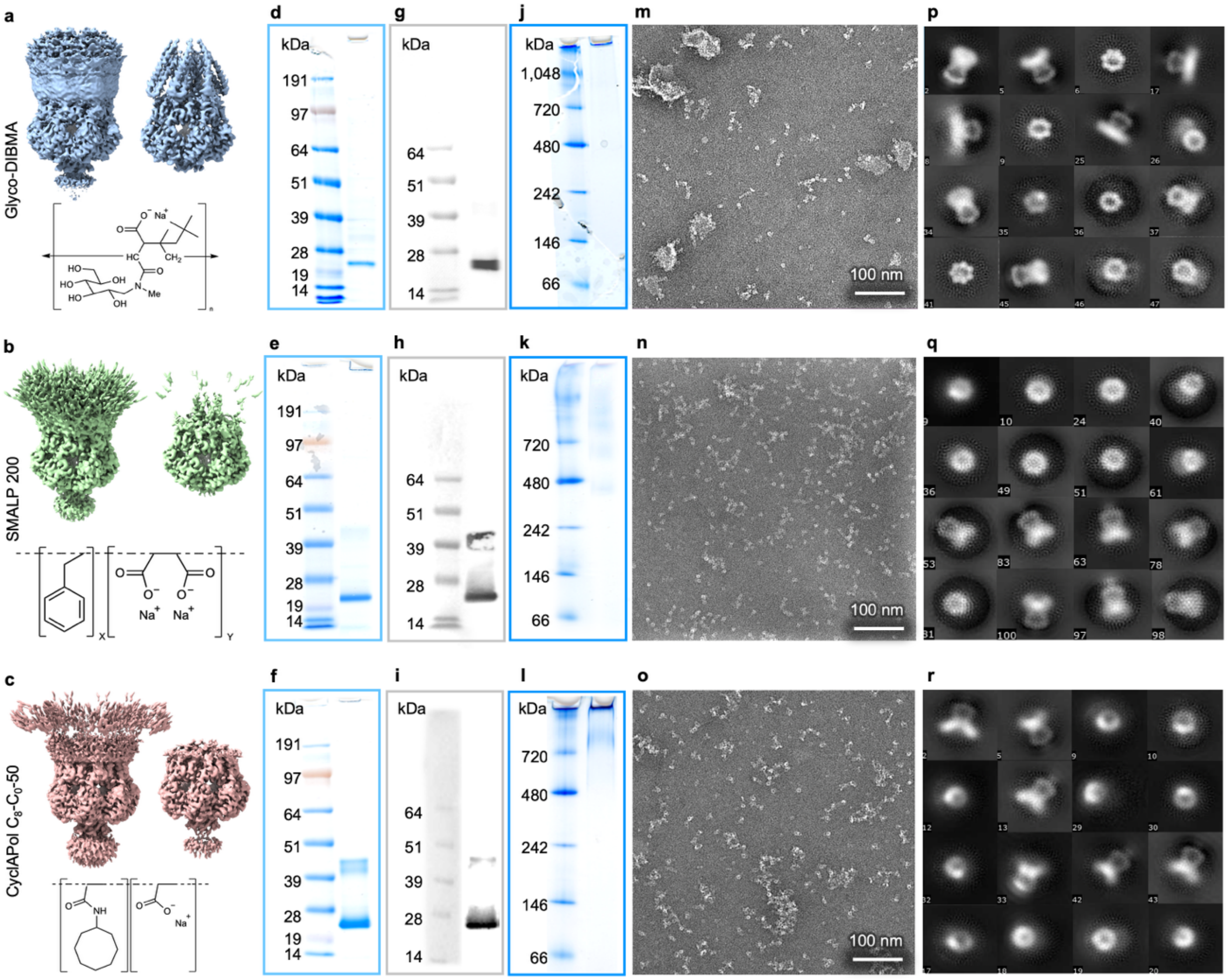


**Extended Data Fig.3: Comparison of electron microscopy and biochemistry of the different polymers.** Top to bottom MscS extracted with 0.5 % Glyco-DIBMA [5], 1% SMALP200 [3], and 0.1% CyclAPol C$_8$-C$_0$-50 [4]. (a-c) 3D reconstructions at varying thresholds, and chemical structures. (d-f) SDS gel indicating the 31 kDa MscS monomer running between the 19 and 28 kDa marker bands. (g-i) Western blot indicating the C-terminally His6-tagged MscS monomer. (j-l) BN-PAGE gel not indicating a nice band at the expected MW of 217 kDa for the MscS heptamer but signal in the gel loading pocket. (m-o) Representative negative staining electron micrograph of MscS extracted with each of the polymers. For 0.5% Glyco-DIBMA (m), there are some isolated single particles as well as small vesicle- or membrane-patch like particles. For both 1% SMALP200 and 0.1% CyclAPol C$_8$-C$_0$-50 (n,o) there are some isolated single particles as well as clusters/aggregation. (p-r) Selected 2D class averages after negative staining EM data collection with a box size of 168 px (~250 Å).

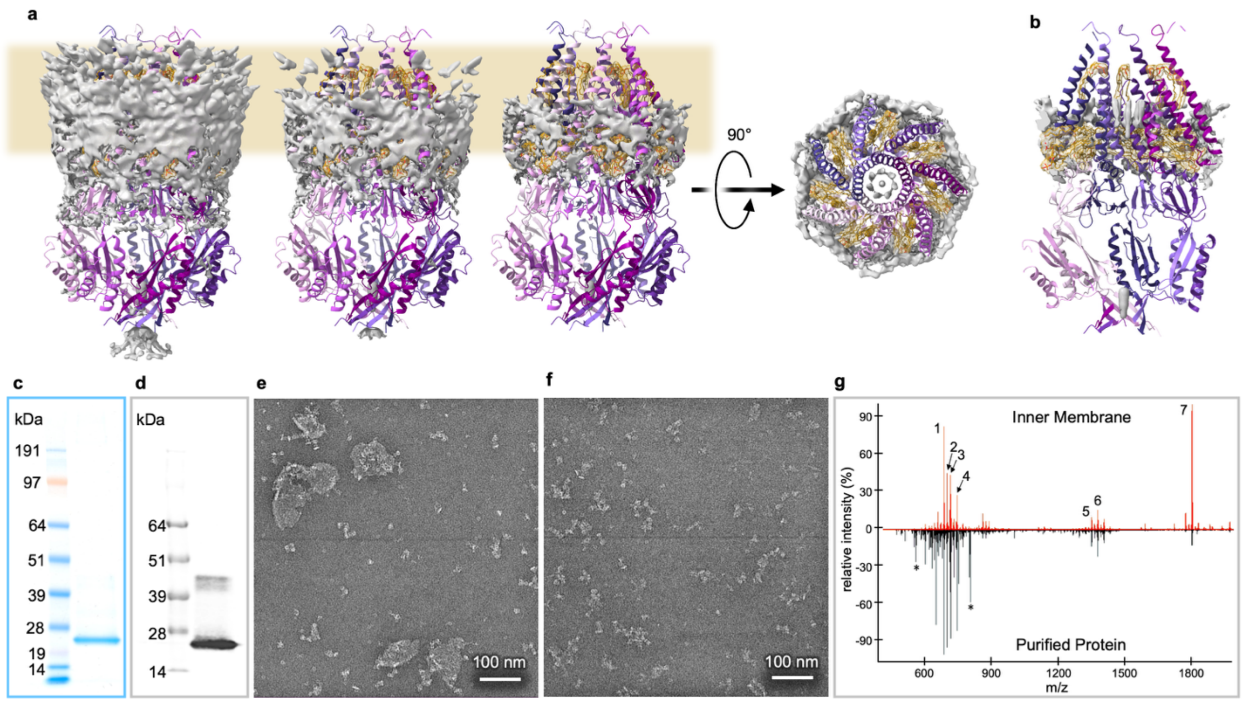


**Extended Data Fig. 4: Extra densities, modeled lipids, and lipid mass spectrometry.** (a) Additional densities not modeled as lipids in C7 symmetry at varying thresholds, side and top views. (b) Side view sliced in half to show unmodeled densities inside the pore. (c-d) SDS gel and Western blot for the sample used for mass spectrometry indicating the C-terminally His6-tagged MscS monomer. (e) Representative negative staining electron micrograph of MscS extracted with Glyco-DIBMA before gel filtration, indicating some isolated single particles as well as small vesicle- or membrane patch-like particles. (f) Representative negative staining electron micrograph after gel filtration showing primarily single particles and lacking the small vesicle- or membrane patch-like particles. (g) Lipid mass spectra for the inner membrane fraction compared to the isolated MscS particles after size exclusion, the numbers show the identified lipids listed in Extended Data Table 2.

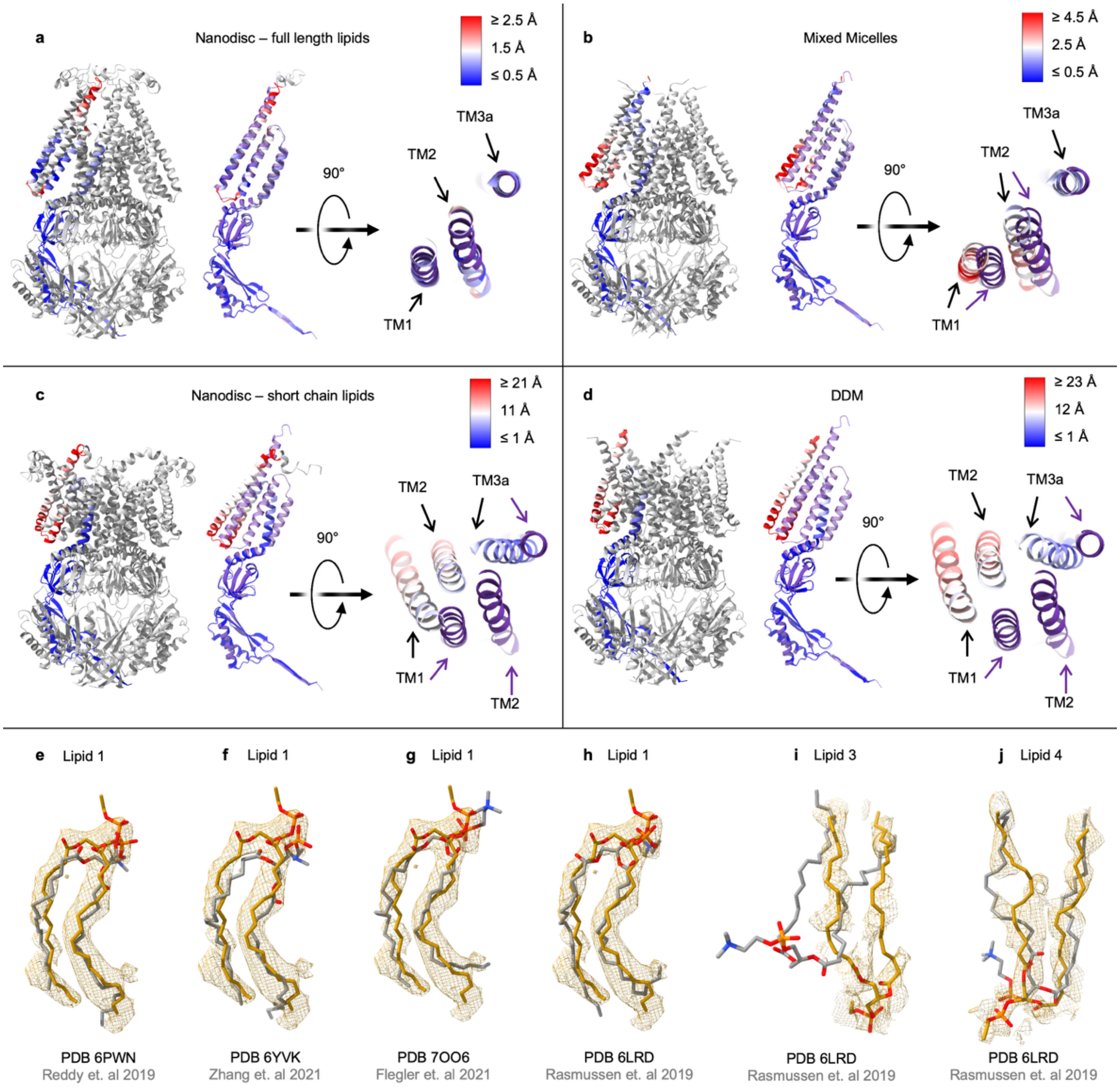


**Extended Data Fig. 5: Structure comparison.** A cartoon representation of MscS with chain A colored by RMSD to the Glyco-DIBMA structure (purple) for (a) nanodiscs with full length lipids (PDB 6PWN [14]), (b) MscS in mixed micelles (PDB 7OO6 [12]), (c) MscS in nanodiscs with short chain lipids (PDB 8DDJ [15]), and (d) MscS in DDM (PDB 7OO0 [12]). Top view comparing the TM helices is on the right. The average RMSD is 0.9 Å, 1.7 Å, 8.6 Å, and 9.6 Å, respectively. A comparison of the lipids resolved in the Glyco-DIBMA MscS structure (gold), shown inside mesh density, with the conformations of previously reported lipids (gray) for Lipid 1 (e-h), Lipid 3 (i), and Lipid 4 (j). The PDB containing the lipid model is listed beneath each lipid comparison [12-14,16].

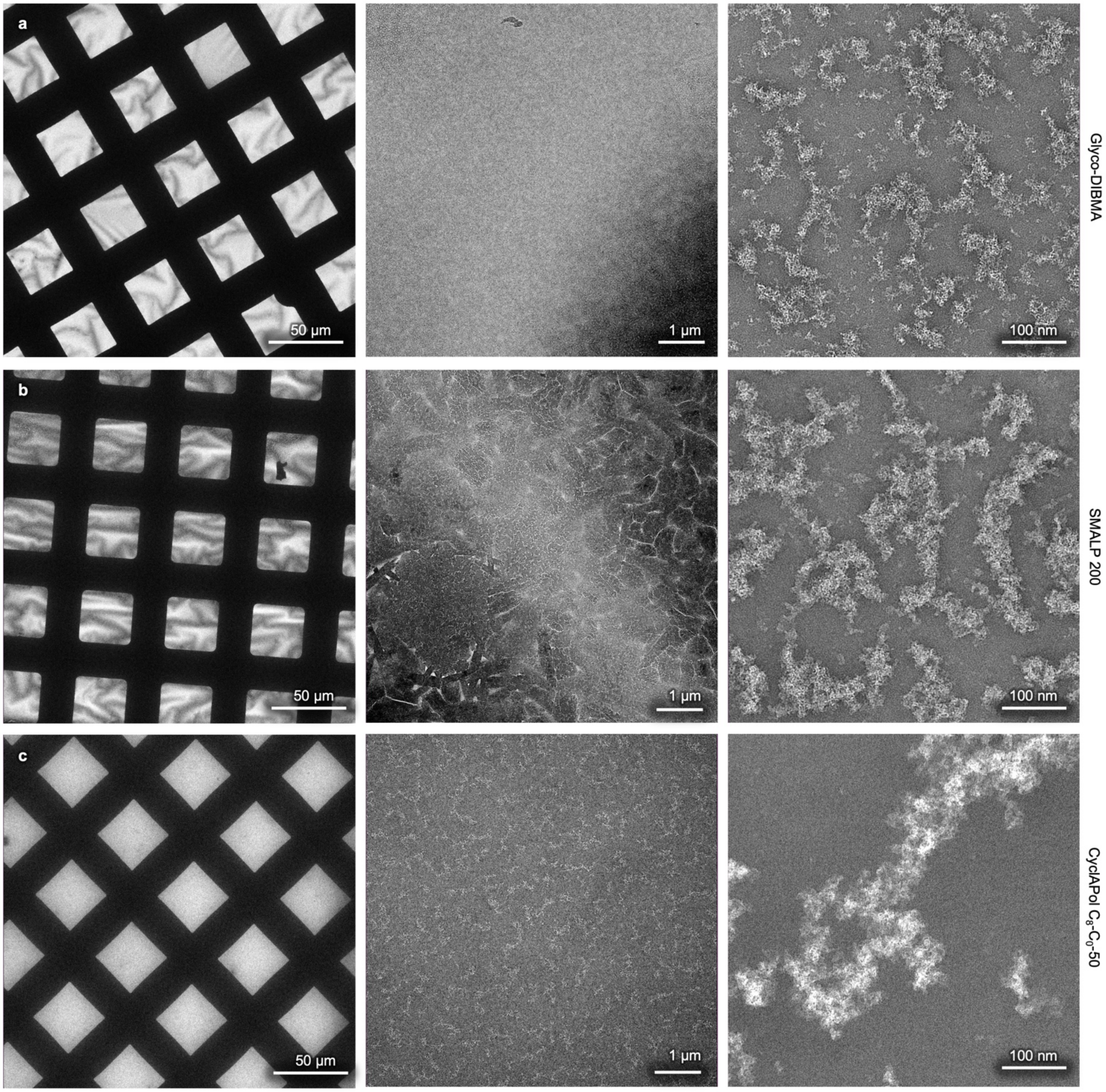


**Extended Data Fig. 6: Negative Staining EM of polymers.** (a) 0.5% Glyco-DIBMA [5] in 20 mM HEPES pH 7.4 Membrane Buffer 1 at low magnification (left), medium magnification (middle), and high magnification (right). (b) 1% SMALP 200 [3] in 20 mM Tris/HCl pH 8.0 Purification Buffer at low magnification (left), medium magnification (middle), and high magnification (right). (c) 0.1% CyclAPol $C_8$-$C_0$-50 [4] in 20 mM Tris/HCl pH 8.0 Purification Buffer at low magnification (left), medium magnification (middle), and high magnification (right).

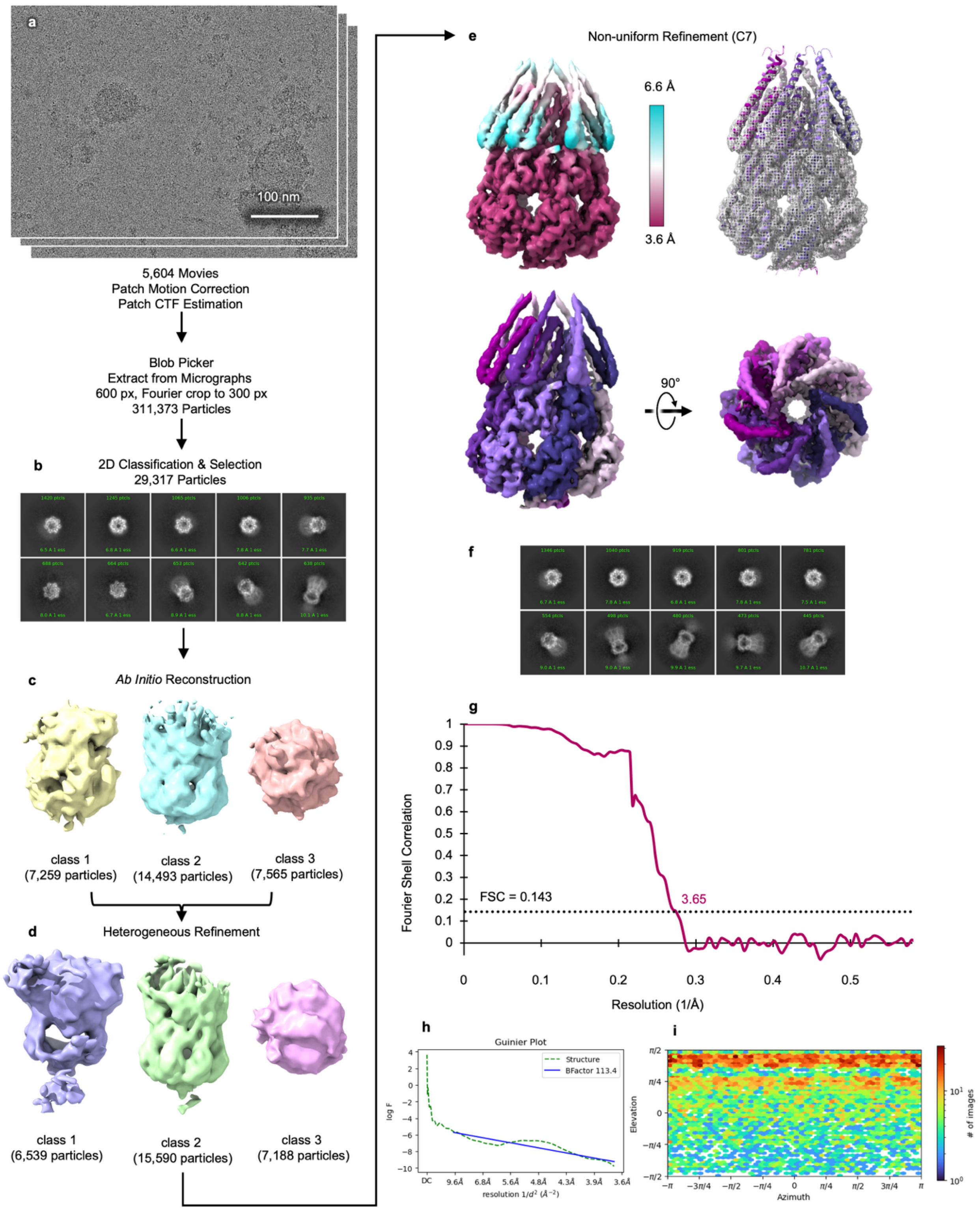


**Extended Data Fig. 7: Glyco-DIBMA #1 cryo-EM workflow.** (a) Representative cryo-EM micrograph. (b) Selected 2D class averages of particles used for *Ab Initio* Reconstruction with a box size of 360 px (~300 Å). (c) *Ab Initio* reconstructions. (d) Heterogeneous Refinement. (e) The final map after non-uniform refinement in C7 symmetry, filtered and colored by local resolution (top left) with the model rigid body fit into the density (top right), and colored by chain (bottom), side view and top view. (f) Representative 2D class averages of particles used for final reconstruction. (g) FSC curve. (h) Guinier Plot. (i) Viewing Direction Distribution Plot.

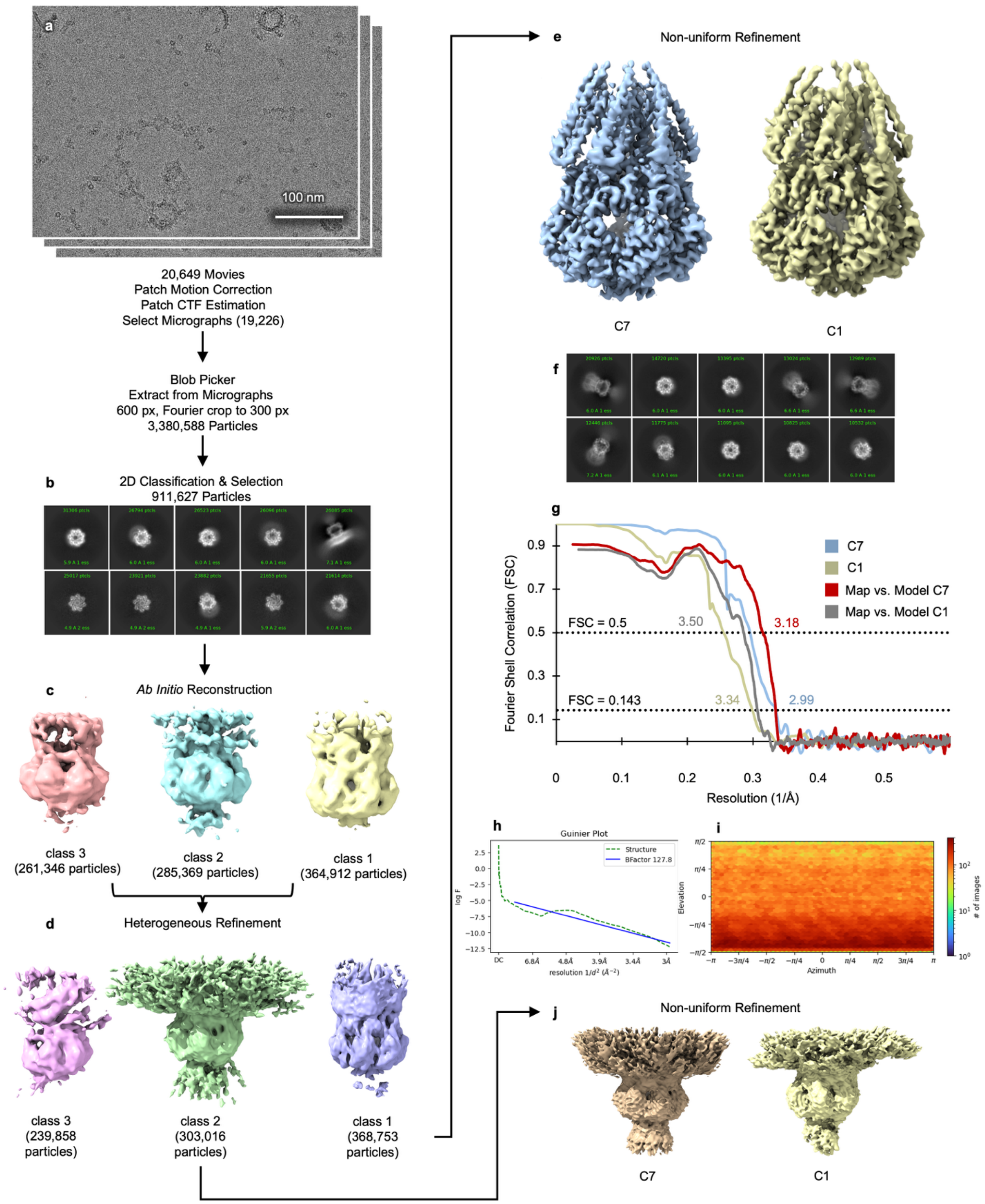


**Extended Data Fig. 8: Glyco-DIBMA #2 cryo-EM workflow.** (a) Representative cryo-EM micrograph. (b) Selected 2D class averages of particles used for *Ab Initio* Reconstruction with a box size of 360 px (~300 Å). (c) *Ab Initio* Reconstructions. (d) Heterogeneous Refinement. (e) Final map after non-uniform refinement in C7 (left) and C1 symmetry (right). (f) Representative 2D class averages of particles used for final reconstruction. (g) FSC curve for C7 (blue), C1 (yellow), and map vs. model in C7 (red) or C1 (gray). (h) Guinier Plot. (i) Viewing Direction Distribution Plot. (j) Non-uniform refinement for Class 2 in d with C7 (left) and C1 symmetry (right).

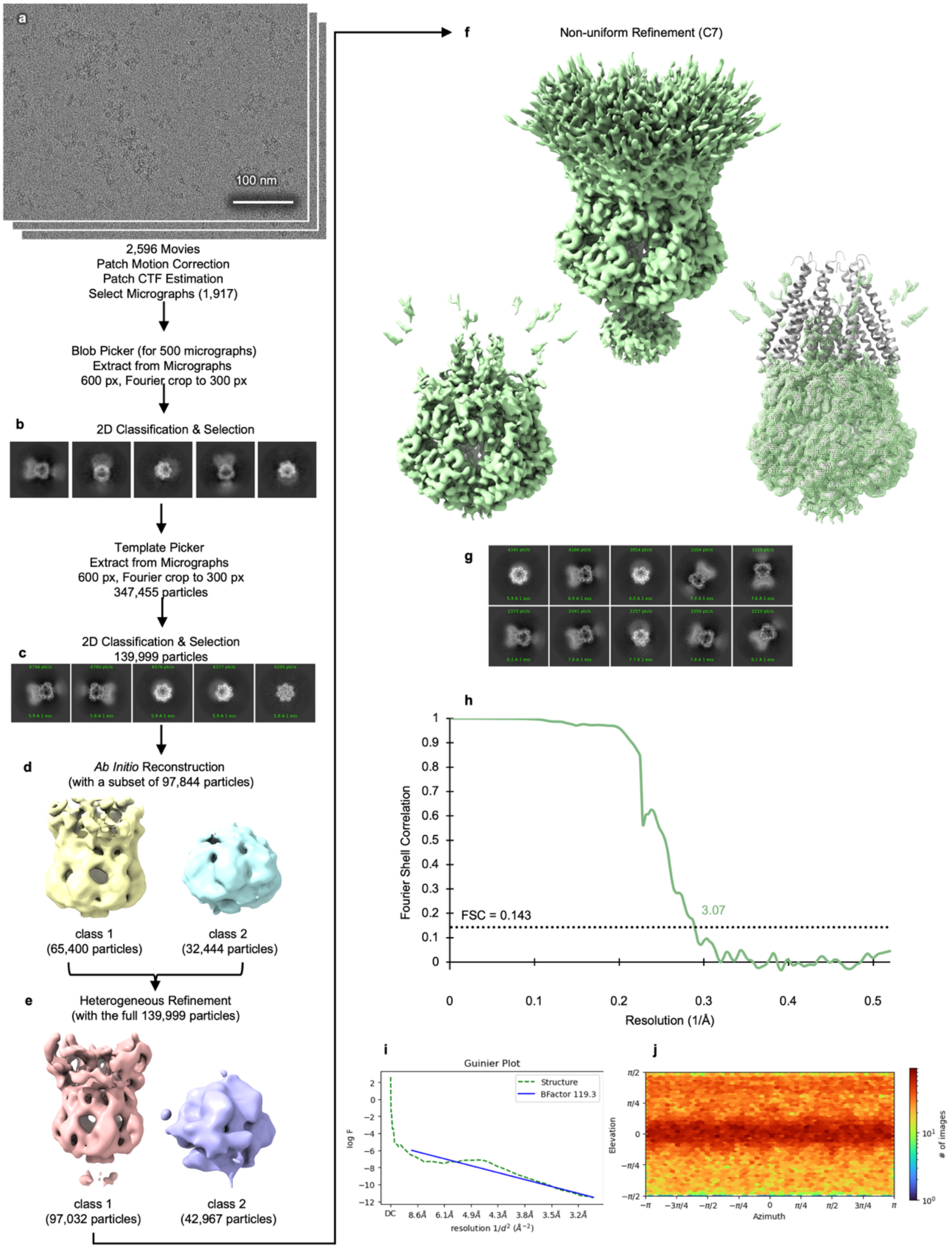


**Extended Data Fig. 9: SMALP 200 #1 cryo-EM workflow.** (a) Representative cryo-EM micrograph. (b) 2D class averages used for template picking, generated from using blob picker on 500 micrographs with a box size of 640 px (~270 Å) Fourier cropped to box size of 320 px. (c) Selected 2D class averages of particles later used for Heterogeneous Refinement (d) *Ab Initio* Reconstructions using a subset of 97,844 particles. (e) Heterogeneous Refinement with the full 139,999 particles. (f) Final map after non-uniform refinement in C7 symmetry (top) shown with a lower threshold (bottom left) and with the model rigid body fit into the density shown in mesh (bottom right). (g) Representative 2D class averages of particles used for final reconstruction. (h) FSC curve. (i) Guinier Plot. (j) Viewing Direction Distribution Plot.

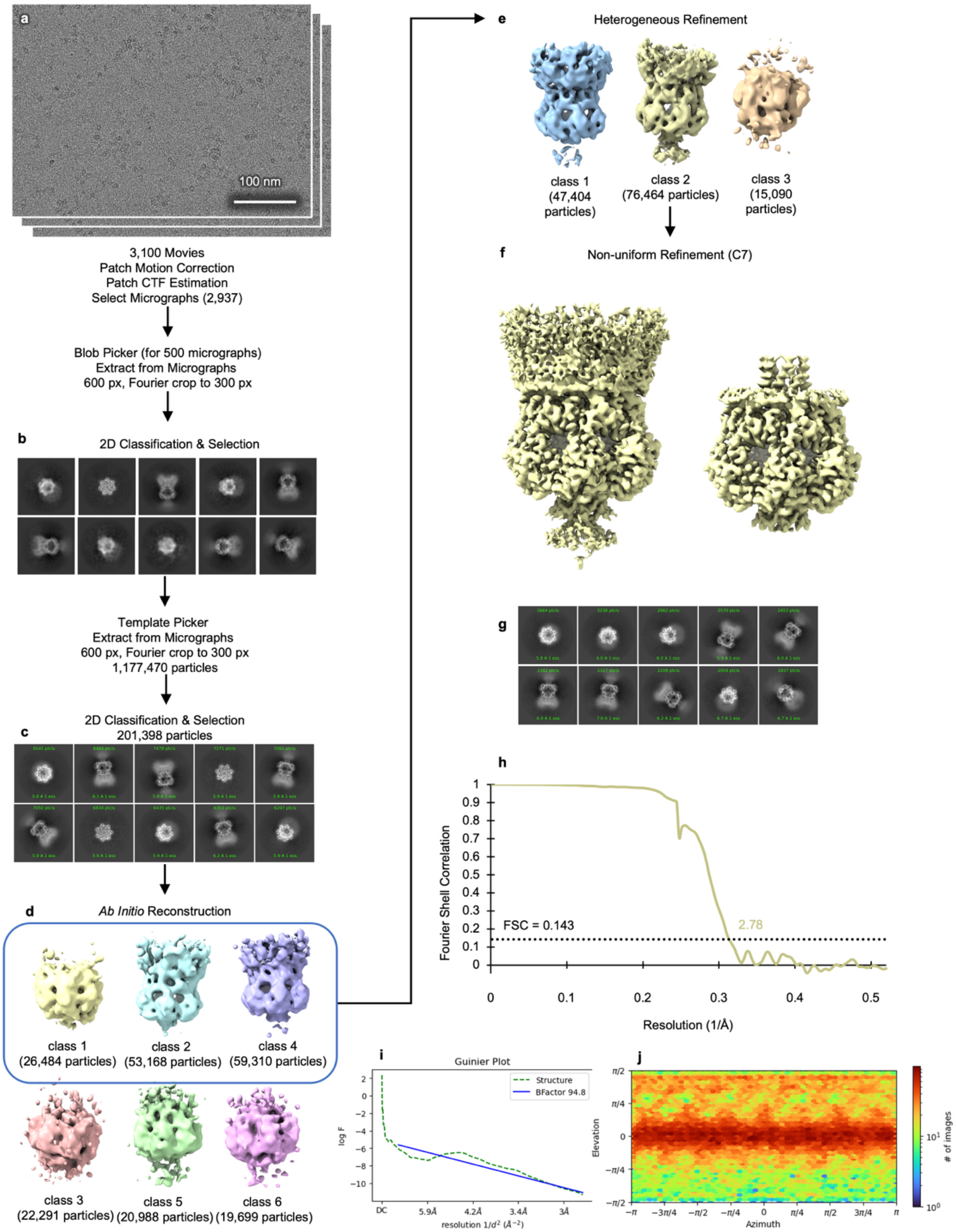


**Extended Data Fig. 10: SMALP 200 #2 cryo-EM workflow.** (a) Representative cryo-EM micrograph. (b) 2D class averages used for template picking, generated from using blob picker on 500 micrographs with a box size of 640 px (~270 Å) Fourier cropped to box size of 320 px. (c) Selected 2D class averages of particles used for *Ab Initio* Reconstruction. (d) *Ab Initio* Reconstructions, particles and maps in the blue box were used for Heterogeneous Refinement (e) Heterogeneous Refinement (f) Final model after non-uniform refinement in C7 symmetry at different thresholds. (g) Representative 2D class averages of particles used for final reconstruction. (h) FSC curve. (i) Guinier Plot. (j) Viewing Direction Distribution Plot.

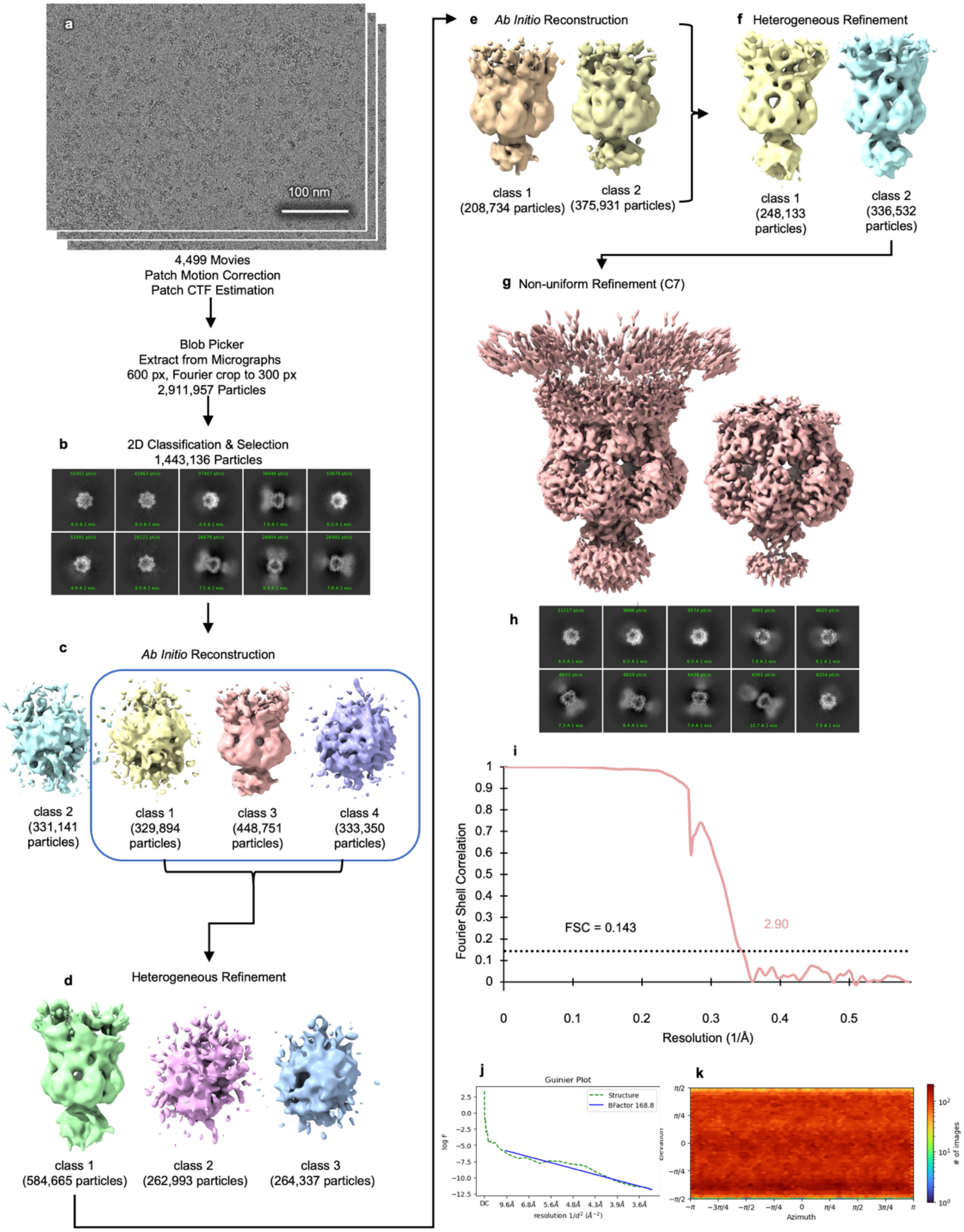


**Extended Data Fig. 11: CyclAPol $C_8$-$C_0$-50 cryo-EM workflow.** (a) Representative cryo-EM micrograph. (b) Selected 2D class averages used for *Ab Initio* Reconstruction with a box size of 360 px (~300 Å). (c) *Ab Initio* Reconstructions, particles and maps in blue box were used for Heterogeneous Refinement. (d) Heterogeneous Refinement, particles from class 1 subjected to another round of *Ab Initio* Reconstruction and divided between two maps. (e) Second round of *Ab Initio* Reconstruction, used for the second round of Heterogeneous Refinement. (f) Second round of Heterogeneous Refinement. (g) Final model after non-uniform refinement in C7 symmetry at different thresholds. (h) Representative 2D class averages of particles used for final reconstruction. (i) FSC curve. (j) Guinier Plot. (k) Viewing Direction Distribution Plot.

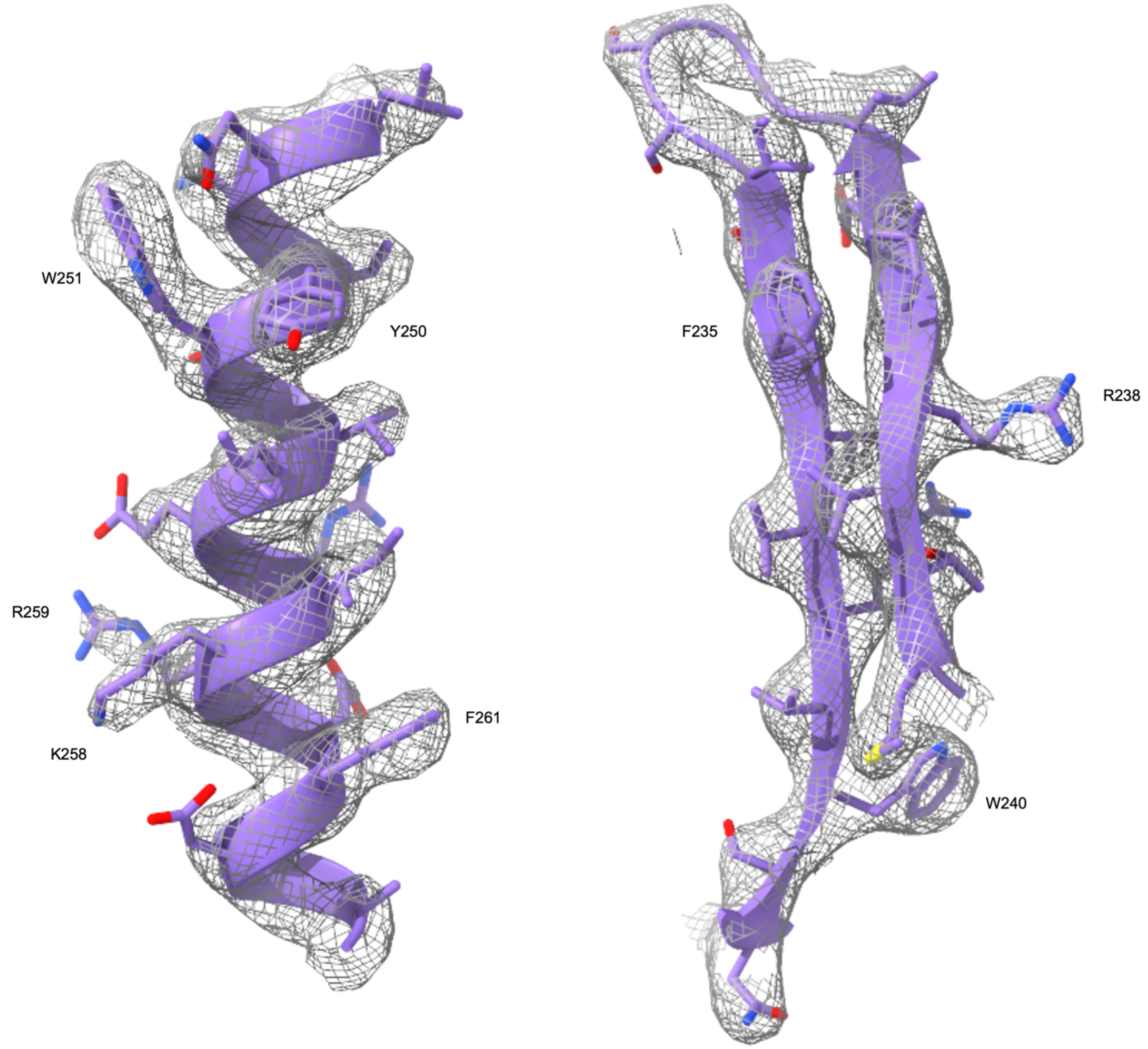


**Extended Data Fig. 12: Quality of the cryo-EM map and fitted model of MscS in Glyco-DIBMA.**
Representative densities of an α-helix (left) and a β-sheet (right) with fitted model in purple in ribbon representation and side chains in sticks.

### Extended Data Tables

**Extended Data Table 1: Protein residues within 5 Å of modeled lipids in Chain A from the static cryo-EM structure.**

| Chain | Lipid 1 | Lipid 2 | Lipid 3 | Lipid 4 |
|---|---|---|---|---|
| A | V29, A33, A36, I37, V40, G41, I44, I77, F80, T81, R88 | I48, V52, L55, R59, L72, V73, G76, F80 | V52, R59, I61, L69, V73, V107, L111, G149, I150, F151 | F68, L69, G104, V107, G108, L111, L115, L118, I150 |
| B | L23, S26, Y27, N30, I31, L82, L86, V89, G90, V91 | Y75, G76, A79, F80, L100, A103, G104, V107 | A103 | T64, V65, A66, D67, F68, A103, A106 |
| C | | V99 | | |
| G | | | M126, F127 | L118, A119, V122, L123, M126, F127 |

**Extended Data Table 2: Values for lipid mass spectrometry analysis.** The ratio of mole fractions of specific lipids in the protein sample isolated in native nanodiscs (S) divided by the total inner membrane fraction (M). The numbers listed in column 1 correspond to the labeled peaks in the spectra shown in Extended Data Fig. 4g. This experimental result was obtained from 9 repeated measurements, conducted across 3 repeated experiments on separate days. Error listed is standard deviation.

| Number | Theoretical $[M\text{-}H]^-$ | Lipids | Chemical formula (Neutral form) | Ratio (S/M) |
|---|---|---|---|---|
| 1 | 688.492 | POPE 32:1 | $C_{37}H_{72}NO_8P$ | 0.99 ± 0.34 |
| 2 | 702.508 | POPE 33:1 | $C_{38}H_{74}NO_8P$ | 2.22 ± 0.21 |
| 3 | 719.487 | POPG 32:1 | $C_{38}H_{73}O_{10}P$ | 2.95 ± 1.51 |
| 4 | 747.518 | POPG 34:1 | $C_{40}H_{77}O_{10}P$ | 2.94 ± 1.58 |
| 5 | 1347.934 | CL 64:2 | $C_{73}H_{138}O_{17}P_2$ | 1.95 ± 1.57 |
| 6 | 1375.965 | CL 66:2 | $C_{75}H_{142}O_{17}P_2$ | 1.91 ± 1.55 |
| 7 | 1796.212 | Lipid A | $C_{94}H_{178}N_2O_{25}P_2$ | 0.44 ± 0.46 |

**Extended Data Table 3: Statistics from MD simulations of lipid contacts.** The probabilities of polar contacts are calculated for lipid phosphate and fatty acid ester oxygens, as well as for the terminal amino groups. The numbers are derived from a 100 ns MD trajectory and represent time-averaged sums for seven subunits.

| | | | **All membrane lipids** | **POPG 301** | **POPG 302** | **POPG 303** | **POPE 304** |
|---|---|---|---|---|---|---|---|
| **R54** | **Lipid oxygens** | **Total** | 14.18 | 0.00 | 0.95 | 0.00 | 0.00 |
| | | **Fatty acid** | 2.79 | 0.00 | 0.94 | 0.00 | 0.00 |
| | | **Phos-phate** | 10.86 | 0.00 | 0.00 | 0.00 | 0.00 |
| **R59** | **Lipid oxygens** | **Total** | 15.68 | 0.00 | 6.17 | 7.84 | 0.01 |
| | | **Fatty acid** | 5.60 | 0.00 | 0.44 | 4.85 | 0.01 |
| | | **Phos-phate** | 9.01 | 0.00 | 5.40 | 2.59 | 0.00 |
| **R74** | **Lipid oxygens** | **Total** | 8.08 | 0.00 | 0.52 | 0.02 | 0.00 |
| | | **Fatty acid** | 0.60 | 0.00 | 0.01 | 0.00 | 0.00 |
| | | **Phos-phate** | 7.01 | 0.00 | 0.44 | 0.00 | 0.00 |
| **R88** | **Lipid oxygens** | **Total** | 12.26 | 12.26 | 0.00 | 0.00 | 0.00 |
| | | **Fatty acid** | 5.87 | 5.87 | 0.00 | 0.00 | 0.00 |
| | | **Phos-phate** | 5.74 | 5.74 | 0.00 | 0.00 | 0.00 |
| **K60** | **Lipid oxygens** | **Total** | 0.07 | 0.00 | 0.00 | 0.06 | 0.00 |
| | | **Fatty acid** | 0.00 | 0.00 | 0.00 | 0.00 | 0.00 |
| | | **Phos-phate** | 0.02 | 0.00 | 0.00 | 0.02 | 0.00 |
| **D67** | **Lipid hydrogens** | **-OH** | 0.00 | 0.00 | 0.00 | 0.00 | 0.00 |
| | | **-NH3** | 3.55 | 0.00 | 1.85 | 0.10 | 1.42 |
| **All protein side chain nitrogens from POLAR but uncharged residues** | **Lipid oxygens** | **Total** | 20.00 | 5.12 | 0.00 | 0.02 | 0.83 |
| | | **Fatty acid** | 10.48 | 4.59 | 0.00 | 0.00 | 0.00 |
| | | **Phos-phate** | 8.47 | 0.53 | 0.00 | 0.02 | 0.83 |
| **All protein side chain oxygens from POLAR but uncharged residues** | **Lipid hydrogens** | **-OH** | 0.95 | 0.00 | 0.00 | 0.00 | 0.00 |
| | | **-NH3** | 2.28 | 0.53 | 0.03 | 0.09 | 0.69 |

**Extended Data Table 4: Activation tension for mutants patched in PB113.** The midpoint ratio of MscS:MscL and the threshold tension were measured in the PB113 (ΔMscS ΔMscK) strain [17] to compare the mutant MscS to the native MscL, whose midpoint tension is 14 mN/m with a midpoint ratio of 0.6. Error listed is standard deviation.

| **Strain** | **$P_{0.5}$ MscS:$P_{0.5}$ MscL** | **$\gamma$ (mN/m)** |
|---|---|---|
| WT | 0.58 | 7.80 |
| R88A (n=3) | 0.57 ±0.04 | 7.74 ±0.59 |
| R59A (n=2) | 0.45 ±0.04 | 5.69 ±0.08 |
| R59S (n=3) | 0.55 ±0.06 | 7.41 ±0.82 |
| K60A (n=3) | 0.66 ±0.11 | 8.96 ±1.46 |
| D67A (n=2) | 0.70 ±0.01 | 9.50 ±0.09 |
| R59S/K60S (n=2) | 0.49 ±0.01 | 6.22 ±0.68 |
| R59S/K60S/D67A (n=3) | 0.44 ±0.05 | 5.97 ±0.61 |

**Extended Data Table 5. Summary of patch clamp data for inactivation and recovery.** The percent inactivation and the inactivation recovery $\tau$ were measured from plasmids expressing MscS in MJF465 (ΔMscL ΔMscS ΔMscK) [18]. Percent inactivation was taken between the initial pulse and the first test pulse in the Inactivation/Recovery Protocol. The recovery from inactivation ($\tau$) is from a fit of the three test pulses to a mono-exponential equation. Error listed is standard deviation.

| **Strain** | **Inactivation (%)** | | **Inactivation Recovery $\tau$ (s)** | |
|---|---|---|---|---|
| | **+ 30 mV** | **- 30 mV** | **+ 30 mV** | **- 30 mV** |
| WT (n=10) | 41 ± 26 | 74 ± 21 | 1.1 ± 0.4 | 1.3 ± 0.5 |
| R88A (n=4) | 26 ± 8 | 43 ± 5 | 2.8 ± 0.5 | 3.5 ± 1.0 |
| R59A (n=3) | 12 ± 8 | 25 ± 22 | N/A | N/A |
| R59S (n=3) | 7 ± 2 | 33 ± 4 | N/A | N/A |
| R59W (n=1) | 15 | 13 | N/A | N/A |
| R59Y (n=1) | 12 | 3 | N/A | N/A |
| D67A (n=7) | 97 ± 5 | 99 ± 1 | 0.7 ± 0.1 | 0.9 ± 0.4 |
| D67N (n=7) | 77 ± 12 | 94 ± 7 | 1.8 ± 0.6 | 2.1 ± 1.0 |
| R59S/K60S (n=5) | 8 ± 8 | 21 ± 22 | N/A | N/A |
| R59S/K60S/D67A (n=3) | 14 ± 5 | 27 ± 22 | N/A | N/A |

**Extended Data Table 6: Cryo-EM data collection parameters and analysis.**

| | Glyco-DIBMA #1 | Glyco-DIBMA #2 | SMALP 200 #1 | SMALP 200 #2 | CyclAPol $C_8$-$C_0$-50 |
|---|---|---|---|---|---|
| **Date of collection** | 10/3/2023 | 11/9/2023 | 5/9/2023 | 5/23/2023 | 10/4/2023 |
| **Protein concentration** | 0.1 mg/mL | 0.3 mg/mL | 0.3 mg/mL | 0.4 mg/mL | 0.7 mg/mL |
| **Sample volume** | 3 µl | 3 µl | 3 µl | 3 µl | 3 µl |
| **Double sample application/blot** | yes | yes | no | no | no |
| **Grid type** | QF R 1.2/1.3 400 Cu mesh + 2 nm C | QF R 1.2/1.3 400 Cu mesh + 2 nm C | QF R 1.2/1.3 400 Cu mesh + 2 nm C | QF R 1.2/1.3 400 Cu mesh + 2 nm C | QF R 1.2/1.3 400 Cu mesh + 2 nm C |
| **Plunge freezer** | Leica EM GP2 | Leica EM GP2 | Leica EM GP2 | Leica EM GP2 | Leica EM GP2 |
| **Blotting time (s)** | 6 | 6 | 6 | 6 | 6 |
| **Temperature (°C)** | 4 | 4 | 4 | 4 | 4 |
| **Humidity (set)** | 95% | 95% | 95% | 95% | 95% |
| **Microscope** | Titan Krios G4 | Titan Krios G4 | Titan Krios G4 | Titan Krios G1 | Titan Krios G4 |
| **Voltage (kV)** | 300 | 300 | 300 | 300 | 300 |
| **Camera** | K3 (non-CDS mode) | K3 (non-CDS mode) | K3 (CDS mode) | K3 (CDS mode) | K3 (non-CDS mode) |
| **Energy filter (slit)** | Yes (20 eV) | Yes (20 eV) | Yes (20 eV) | Yes (20 eV) | Yes (20 eV) |
| **Cs corrector** | no | no | no | no | no |
| **Objective aperture** | no | no | no | 100 µm | no |
| **Magnification** | 105,000x | 105,000x | 105,000x | 105,000x | 105,000x |
| **Physical pixel size (Å/px)** | 0.8469 | 0.8469 | 0.8469 (0.425 super-res) | 0.83 (0.415 super-res) | 0.8469 |
| **Electron exposure ($e^-$/Å$^2$)** | 50 | 50 | 50 | 50 | 50 |
| **Number of movie frames** | 60 | 60 | 60 | 50 | 60 |
| **Dose rate ($e^-$/px/s)** | 15 | 15 | 10 | 9 | 15 |
| **Defocus (µm)** | -0.8 to -1.7 | -0.8 to -1.7 | -0.8 to -1.7 | -0.8 to -1.8 | -0.8 to -1.7 |
| **Number of total micrographs** | 5,604 | 20,649 | 2,597 | 3,100 | 4,654 |
| **Number of selected micrographs** | 5,604 | 19,226 | 1,917 | 2,937 | 4,654 |
| **Number of particles picked** | 311,373 | 3,380,588 | 47,038 | 1,177,470 | 2,911,957 |

**Extended Data Table 7: Cryo-EM map and model analysis.**

| | **Glyco-DIBMA #2 in C7**<br>**EMD-71088**<br>**PDB ID: 7P0N** | **Glyco-DIBMA #2 in C1**<br>**EMD-71089**<br>**PDB ID: 7P0O** |
|---|---|---|
| **Data processing** | | |
| Final number of particles | 368,753 | 368,753 |
| Final pixel size used for final maps (Å/px) | 0.8469 | 0.8469 |
| Symmetry imposed | C7 | C1 |
| Resolution of map (Å) | 2.99 (2.5-4.5) | 3.34 (2.9-6.3) |
| FSC threshold | 0.143 | 0.143 |
| *B* factor for map ($Å^2$) | 127.8 | 97.2 |
| **Model composition** | | |
| Chains | 7 | 7 |
| Non-hydrogen atoms | 15,499 | 14,014 |
| Protein residues | 1,855 | 1,855 |
| Water | 91 | 0 |
| Ligands | 28 | 0 |
| *B* factors ($Å^2$) | | |
| Protein | 79.46 | 96.57 |
| Ligand | 151.08 | N/A |
| **Refinement and validation** | | |
| Initial model used (PDB code) | 7OO6 | 7OO6 |
| Model resolution (Å) | 3.17 | 3.50 |
| FSC threshold | 0.5 | 0.5 |
| R.m.s. deviations | | |
| Bond lengths (Å) | 0.004 | 0.003 |
| Bond angles (°) | 0.527 | 0.557 |
| Validation | | |
| MolProbity score | 1.01 | 1.61 |
| Clashscore | 2.30 | 5.46 |
| Poor rotamers (%) | 0.48 | 1.91 |
| Ramachandran plot | | |
| Favored (%) | 98.10 | 97.50 |
| Allowed (%) | 1.90 | 2.50 |
| Disallowed (%) | 0.00 | 0.00 |

**Primers (5’-3’)**

R59A F: GATGATCTCCgcgAAAATCGATGCCACTGTTGCTG
R59S F: GATGATCTCCagcAAAATCGATGCCACTGTTG
R59W F: GATGATCTCCtggAAAATCGATGCC
R59Y F: GATGATCTCCtatAAAATCGATGCCACTGTTGC
R59A/K60A F: GATGATCTCCgcggcgATCGATGCCACTGTTGCTGATTTTC
R59S/K60S F: GATGATCTCCagcagcATCGATGCCACTGTTGCTG
R59 R: AGGCGATTCACCGCGTTG
K60A F: GATCTCCCGTgcgATCGATGCCACTG
K60S F: GATCTCCCGTagcATCGATGCCACTG
K60 R: ATCAGGCGATTCACCGCG
D67A F: CACTGTTGCTgcgTTTCTTTCTGCATTAG
D67N F: CACTGTTGCTaacTTTCTTTCTG
D67 R: GCATCGATTTTACGGGAG

### Supplementary Methods

*Protein Validation using SDS-PAGE, Western Blot, and BN-PAGE*

Protein fractions were resolved and analyzed using SDS-PAGE, Western Blot, and BN-PAGE. For SDS-PAGE samples were prepared with 4X Bolt LDS Sample Buffer (Invitrogen) and incubated at 80°C for 10 minutes before loading into 4-12% Bis-Tris gel (Invitrogen) with NuPAGE MOPS SDS Buffer (Invitrogen) and running at 80 V for 15 minutes followed by 200 V for 45 minutes on ice. SDS-gel was fixed and then stained with Colloidal Blue Stain (Invitrogen) overnight and de-stained in water and scanned the following day using Espson Perfection V850 Pro. For Western Blot, SDS-PAGE gel was transferred to PVDF membrane at 30 V for 45 min in NuPAGE Transfer Buffer (Invitrogen) and blocked with 4% BSA in TTBS (100 mM Tris/HCl pH 7.5, 150 mM NaCl, 0.1% Tween 20) for 1 hour, incubated with 0.1 µg/mL Anti-6X His tag antibody (abcam) in 2% BSA in TTBS for 1 hour, washed with TTBS and imaged with ChemiDoc (BioRad) using WesternBright (advansta). Note that the MscS monomer is 31 kDa but migrates faster than expected for its size. This is not unusual, as membrane proteins have been previously shown to have anomalous gel mobility [19]. For BN-PAGE samples were prepared with NativePAGE Sample Buffer and 0.125% NativePAGE G-250 Sample Additive (Invitrogen) and loaded into a 4-16% Bis-Tris gel (Invitrogen) with NativePAGE Running Buffer and NativePAGE Cathode Buffer in the inner chamber (Invitrogen) before running at 150 V for 1 hour followed by 250 V for 1 hour on ice. Gel was fixed in 40% ethanol, 10% acetic acid microwaved for 1 minute and then stained with Colloidal Blue Stain (Invitrogen) overnight and de-stained in water and scanned the following day. Gel images and Western Blots for each purification can be found in Extended Data Fig. 3d-l.

*SMALP 200 Extraction and Affinity Purification*

*E. coli* inner membranes were resuspended to 20 mg/mL in Membrane Buffer 2 (50 mM Tris/HCl pH 8.0, 500 mM NaCl, 10% glycerol [v:v]) and homogenized before the addition of 1% [w:v] of SMALP 200 [3] (Orbiscope) and allowed to extract for 2 hrs at room temperature on a tube revolver. SMALP 300 was also attempted but did not successfully extract. The insoluble fraction was then pelleted by ultracentrifugation at 25,000 rpm using Beckman SW28 rotor (112,400 x g) for 35 min at 4ºC. The supernatant was then diluted 1:4 to improve binding efficiency and incubated overnight on a tube revolver at 4ºC with TALON Cobalt Resin (Takara Bio). The next day the resin was washed with 10 column volumes of Membrane Buffer 2 and 10 column volumes of Membrane Buffer 2 with 20 mM imidazole before elution with Membrane Buffer 2 with 300 mM imidazole. Elution fractions were then dialyzed with Purification Buffer (20 mM Tris/HCl pH 8.0, 250 mM NaCl, 5% glycerol [v:v]) overnight to remove imidazole and concentrated the following day using a 100 kDa ultrafiltration membrane (Amicon) to desired concentration for cryo-EM grid preparation. Protein fractions were resolved and analyzed using SDS-PAGE, Western Blot, and BN-PAGE. Gel images and Western Blots can be found in Extended Data Fig. 3e,h,k.

*CyclAPol $C_8$-$C_0$-50 Extraction and Affinity Purification*

*E. coli* inner membranes were resuspended to 10 mg/mL in Purification Buffer and homogenized before the addition of 0.1% [w:v] of CyclAPol $C_8$-$C_0$-50 [4] and allowed to extract for 2 hrs at room temperature on a tube revolver. CyclAPol $C_8$-$C_0$-50 was provided by Manuela Zoonens (Université de Paris and Institut de Biologie Physico-Chimique, Paris, France) and is now commercially available as Ultrasolute Amphipol 18 (Cube Biotech). The insoluble fraction was then pelleted by ultracentrifugation at 25,000 rpm using Beckman SW28 rotor (112,400 x g) for 35 min at 4°C and the supernatant was incubated overnight on a tube revolver at 4°C with TALON Cobalt Resin (Takara Bio). The next day the resin was washed with 10 column volumes of Purification Buffer and 10 column volumes of Purification Buffer with 10 mM imidazole before elution using Purification Buffer with 300 mM imidazole. Elution fractions were then dialyzed with Purification Buffer overnight to remove imidazole and concentrated the following day using a 100 kDa ultrafiltration membrane (Amicon) to desired concentration for cryo-EM grid preparation. Protein fractions were resolved and analyzed using SDS-PAGE, Western Blot, and BN-PAGE. Gel images and Western Blots can be found in Extended Data Fig. 3f,i,l.

*Negative Staining EM of Polymers in Buffer*

3 μL of polymer sample of 0.5%, 1%, and 0.1% for Glyco-DIBMA, SMALP 200, and CyclAPol $C_8$-$C_0$-50, in Membrane Buffer, Membrane Buffer 2, and Purification Buffer respectively was applied to a glow discharged carbon coated 400 square mesh copper grid (CF400-CU, EMS) and incubated for 1 min at room temperature. The grid was then blotted by filter paper (Whatman Grade 2) and washed once with 3 μL of Nano-W negative staining solution (Nanoprobes) followed by incubation with 3 μL of Nano-W for 1 min. The grid was blotted to remove excess staining solution and air-dried by waving for an additional minute. Images were recorded using an FEI Tecnai T20 TEM operated at 200 kV with a direct electron detector K2 Summit (Gatan Inc) (See Extended Data Fig. 6).

*Negative Staining Data Collection for SMALP 200 and CyclAPol $C_8$-$C_0$-50*

Data was collected using SerialEM [20] at a nominal magnification of 25,000x with a pixel size of 1.479 Å/px, dose rate of approximately 7 $e^-$/px/s with 0.2 s frames over 10 s, and a defocus range between -1.3 and -2 μm. A total of 645 and 509 images were collected for SMALP 200 and CyclAPol $C_8$-$C_0$-50, respectively. Bsoft 2.2.0 [21] was used for unpacking the image stacks. Image processing was performed using cisTEM v1.0.0. [22]. 237,492 and 158,613 particles were picked from the micrographs after CTF estimation [23] for SMALP 200 and CyclAPol $C_8$-$C_0$-50, respectively, using a maximum radius of 60 Å and a characteristic radius of 40 Å with a threshold of 5. Particles were extracted with a box size of 168 px (~250 Å) and analyzed using 2D classification (Extended Data Fig. 3p-r).
